# Nicotinic acetylcholine receptor signaling maintains epithelial barrier integrity

**DOI:** 10.1101/2023.02.09.527809

**Authors:** Nadja S. Katheder, Kristen C. Browder, Diana Chang, Ann De Mazière, Pekka Kujala, Suzanne van Dijk, Judith Klumperman, Zijuan Lai, Dewakar Sangaraju, Heinrich Jasper

## Abstract

Disruption of epithelial barriers is a common disease manifestation in chronic degenerative diseases of the airways, lung and intestine. Extensive human genetic studies have identified risk loci in such diseases, including in chronic obstructive pulmonary disease (COPD) and inflammatory bowel diseases (IBD). The genes associated with these loci have not fully been determined, and functional characterization of such genes requires extensive studies in model organisms. Here, we report the results of a screen in *Drosophila melanogaster* that allowed for rapid identification, validation and prioritization of COPD risk genes that were selected based on risk loci identified in human genome-wide association studies (GWAS) studies. Using intestinal barrier dysfunction in flies as a readout, our results validate the impact of candidate gene perturbations on epithelial barrier function in 56% of the cases, resulting in a prioritized target gene list. We further report the functional characterization in flies of one family of these genes, encoding for nicotinic acetylcholine receptor subunits (nAchR). We find that nAchR signaling in enterocytes of the fly gut promotes epithelial barrier function and epithelial homeostasis by regulating the production of the peritrophic matrix. Our findings identify COPD associated genes critical for epithelial barrier maintenance, and provide insight into the role of epithelial nAchR signaling for homeostasis.

## Introduction

Barrier epithelia such as the skin, linings of the gastrointestinal and urogenital tracts and the airways play a critical role in maintaining a strict separation of external and internal environments, yet also enable the exchange of gases, water, nutrients and immune mediators. They serve as a first layer of defense against external insults and possess remarkable regenerative capacity that declines with age (Jasper, 2020). In *Drosophila*, loss of intestinal barrier function is accompanied by commensal dysbiosis and inflammation and reliably predicts impending organismal death (Rera et al., 2012). Similarly, increased barrier permeability and changes in microbiome composition and abundance have been reported in various human diseases, such as inflammatory bowel disease and chronic obstructive pulmonary disease (COPD) (Raftery et al., 2020). COPD is a major contributor to global morbidity and mortality and is characterized by an obstructed airflow resulting in shortness of breath upon exertion. At the tissue level, lungs of COPD patients display chronic inflammation, extensive cellular remodeling and barrier dysfunction (Aghapour et al., 2018; Barnes, 2019; Carlier et al., 2021).

While smoking or exposure to environmental air pollutants remain major risk factors, many COPD patients are non-smokers, suggesting a genetic component contributing to disease susceptibility (Aghapour et al., 2022; Barnes, 2019). Several GWAS studies have been performed in which risk loci for incidence of COPD have been identified (Hobbs et al., 2017; Pillai et al., 2009; Sakornsakolpat et al., 2019). One of the most well-known risk loci is located near the nicotinic acetylcholine receptor CHRNA3/5 genes and has also been associated with increased nicotine dependence and smoking behavior, and lung cancer (Amos et al., 2008; Carlier et al., 2021; Cui et al., 2014; Hobbs et al., 2017; Hung et al., 2008; Pillai et al., 2009; Wilk et al., 2012). Recent work has demonstrated a role for CHRNA5 in the formation of COPD-like lesions in the respiratory epithelium independently of cigarette smoke, suggesting a direct involvement of nAchRs in shaping epithelial integrity (Routhier et al., 2021). The endogenous ligand of nAchR, acetylcholine (Ach), is a classic neurotransmitter synthesized by Choline Acetyltransferase (ChAT) in cholinergic neurons, as well as in immune cells and epithelial cells, such as brush/tuft cells (Kummer & Krasteva-Christ, 2014; Wessler & Kirkpatrick, 2008). Such cells orchestrate type 2 inflammatory responses (O’Leary et al., 2019; Sell et al., 2021), mucociliary clearance (Perniss et al., 2020) and limit biliary inflammation (O’Leary et al., 2022; O’Leary et al., 2019). How Ach influences homeostasis of barrier epithelia and how disease-associated nAchR variants perturb epithelial function remains mostly unclear.

Overall, experimental evidence for the involvement of specific genes associated with the COPD risk loci identified in these studies is mostly lacking, and will be essential for the development of therapeutic strategies targeting novel pathways. Here, we have used the *Drosophila* midgut as a genetically accessible model for epithelial barrier homeostasis to interrogate genes predicted to be involved in COPD based on GWAS studies. The *Drosophila* intestine is lined by a pseudostratified epithelium consisting of enterocytes (ECs) and enteroendocrine cells (EEs) that are regenerated from a basal population of intestinal stem cells (ISCs) (Miguel-Aliaga et al., 2018). In its structure, cell composition and molecular regulation of regenerative processes, the fly intestinal epithelium resembles mammalian airway epithelia (Biteau et al., 2011).

Under stress conditions, in response to enteropathogen infection, as well as during normal aging, the fly intestinal epithelium loses its barrier function and exhibits stem cell hyperplasia and commensal dysbiosis (Jasper, 2020). These phenotypes recapitulate changes observed in airway epithelia of COPD patients and can thus be used as a model for pathophysiological changes occurring in this disease (Carlier et al., 2021; Raftery et al., 2020).

To assess the role of candidate genes associated with risk alleles in COPD GWAS studies in the maintenance of barrier epithelia integrity, we performed an RNA interference screen perturbing their *Drosophila* orthologues systemically and quantifying the impact of these perturbations on intestinal barrier function. Several of the candidate genes identified in this screen as required for barrier integrity encode for subunits of the nicotinic acetylcholine receptor (nAchR).

In the fly intestine, we find that ChAT is expressed by a subset of enteroendocrine cells and that enterocyte-specific expression of nAchR is required for barrier integrity by stimulating chitin release and ensuring maintenance of the peritrophic matrix, a chitinous structure protecting the epithelium from luminal insults. In ECs, Ach is required for the expression of Syt4, a critical regulator of exocytosis (Yoshihara et al., 2005; Zhang et al., 2011) which is required for the maintenance of PM structure and epithelial barrier function. Our data illustrate the usefulness of *Drosophila* as a model for prioritization of potential disease genes identified in GWAS studies, and identify nAchR signaling as a critical mediator of epithelial homeostasis in barrier epithelia.

## Results

### A genetic screen assessing the role of COPD candidate genes in barrier function

To obtain a curated candidate gene list for COPD, we assigned candidate genes to COPD risk loci (Hobbs et al., 2017) using a combination of expression quantitative trait loci, coding annotation and distance-based metrics (see Methods, Table S1A). *Drosophila* orthologs were identified with the DRSC integrative ortholog prediction tool (DIOPT (Hu et al., 2011)) and corresponding hits with the highest DIOPT score were selected, resulting in a total of 33 *Drosophila* genes screened initially (Fig. 1A).

**Figure 1:**
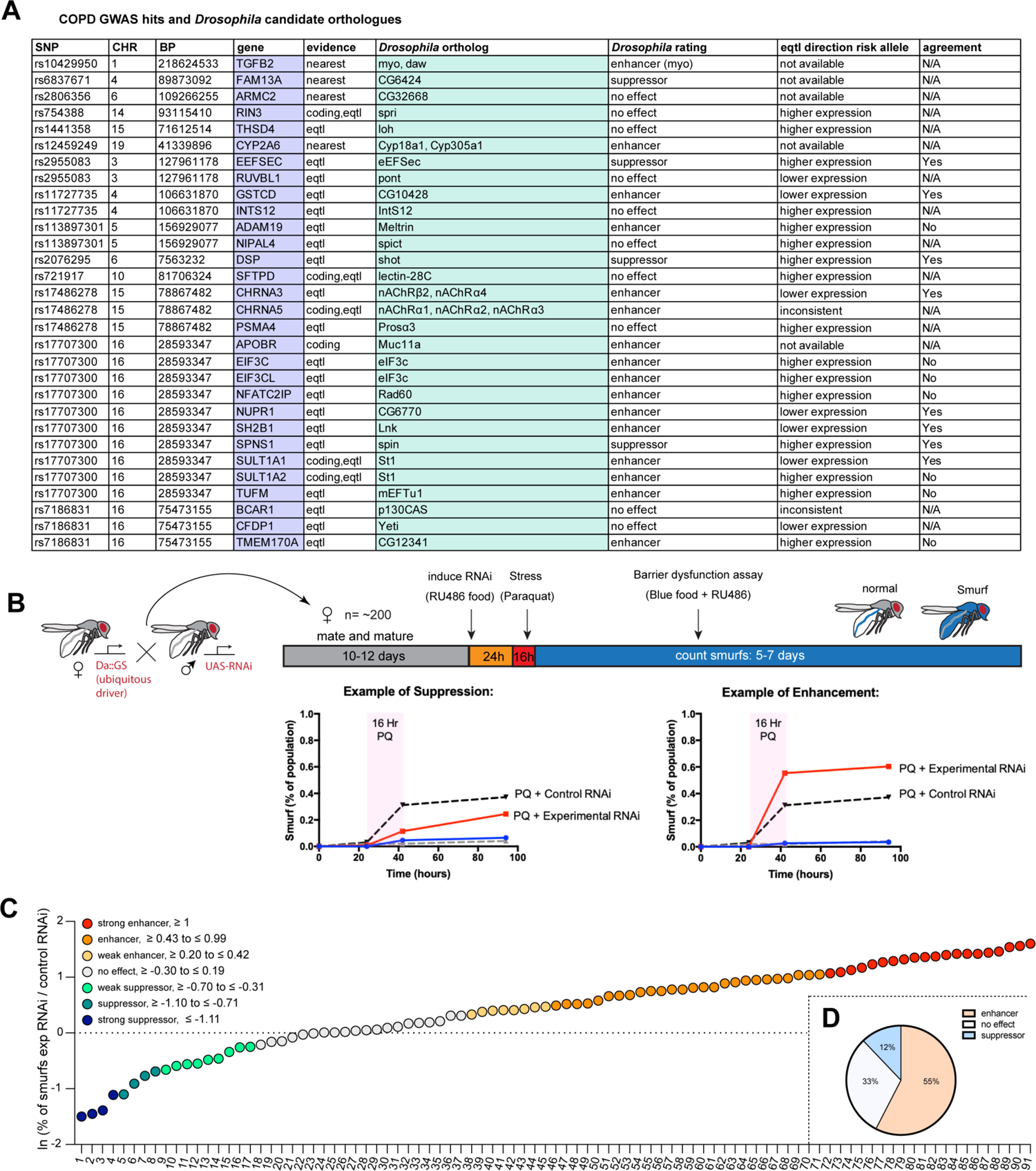
A *Drosophila* screen for COPD-associated candidate genes. A) List of human candidate genes for genetic loci associated with COPD risk and their *Drosophila* orthologs. An overall rating was assigned to the *Drosophila* genes based on the detailed results of the individual RNAi lines included in the screen: Genes exacerbating barrier dysfunction upon depletion were categorized as enhancers, while genes whose depletion improved barrier function were rated as suppressors of barrier dysfunction (Table S1B). When available, the human risk allele expression data is compared to the results of the *Drosophila* screen (agreement column). (SNP, Single nucleotide polymorphism; CHR, chromosome; BP, base pair number; eqtl, expression quantitative trait loci). B) Experimental design of intestinal barrier function screen. Flies carrying the ubiquitous driver Da::GS were crossed to RNAi lines targeting candidate genes. The female offspring was aged for 10-12 days before induction of RNAi expression by RU-486 for 24h on blue food. Flies were challenged with sucrose alone (mock) or 25mM Paraquat for 16h overnight and then placed back on blue food. Blue flies with a defective intestinal barrier(“smurfs”) were counted daily for 5-7 days. C) Ranking of screened RNAi lines based on the natural logarithm (ln) of the ratio between the proportion of smurfs after candidate gene knockdown and luciferase RNAi control. Each number corresponds to an RNAi line listed in Table S1B. Cut offs for the different categories are indicated. D) Summary of screen results based on broad categorization as enhancer, suppressor or no effect. If several RNAi lines targeting the same gene unanimously had no effect, the gene was rated “no effect, conclusive”, while inconsistent results were rated “no effect, inconclusive”. For details see Table S1B.

We perturbed these genes systemically by RNA interference (RNAi) in an inducible fashion using the ubiquitous RU486-inducible Gal4 driver Da::GeneSwitch (Da::GS) and scored epithelial barrier dysfunction in homeostatic and stress conditions using the “smurf assay” (Rera et al., 2011). In this approach flies are fed food containing a non-absorbable blue food dye. If the intestinal epithelial barrier is compromised, the dye leaks into the open circulatory system and gives the fly a blue appearance reminiscent of the popular blue cartoon characters. Where available, a minimum of 2 different RNAi lines per gene were included (Table S1B). Female flies carrying the driver and RNAi construct were allowed to mature and mate for 10-12 days before being placed on blue food with RU-486 to induce knockdown for 24h. Since COPD is strongly associated with environmental stress, we then challenged flies with paraquat (N, N’-dimethyl-4,4’-bipyridinium dichloride), a herbicide known to inflict oxidative stress and damage to the fly gut comparable to the effects of cigarette smoke on the lung epithelium (Biteau et al., 2008; Caliri et al., 2021). After 16h paraquat challenge the flies were moved back to blue food containing RU486 and smurf numbers were recorded over the span of about a week (Fig 1B). We generated a “barrier dysfunction index” for every RNAi line by calculating the natural logarithm (ln) of the ratio of peak smurf percentage between RNAi line and control knockdown and plotted individual RNAi lines accordingly. A positive index implies an enhancement of barrier dysfunction after depletion, while a negative index suggests rescue of barrier integrity after depletion (Fig. 1C). Based on the outcomes of individual RNAi knockdowns, we assigned an overall rating for each candidate gene (Table S1B). We found that disruption of 18 genes (∼55%) resulted in enhancement (e.g. these genes were necessary for barrier integrity), while disruption of 4 genes (12%) resulted in suppression of the barrier dysfunction. The remaining 11 genes did not display any effect on barrier function (Fig. 1D). Out of the 16 of *Drosophila* hits where eqtl data was available for the corresponding human gene, 9 were consistent with the direction of the effect inferred from the association of the COPD risk allele with gene expression (56%, Fig. 1A, Table S1B).

### nAchR subunit expression in ECs is required for barrier function

Our initial screen identified disruption of 5 nAchR subunits as a strong enhancers of barrier dysfunction. Ubiquitous knockdown of various nAchR subunits with Da::GS lead to mild barrier dysfunction under homeostatic conditions, and greatly enhanced barrier dysfunction after paraquat challenge (Fig. 2A, Fig. S1A), suggesting a sensitization of the epithelium to stress.

**Figure 2:**
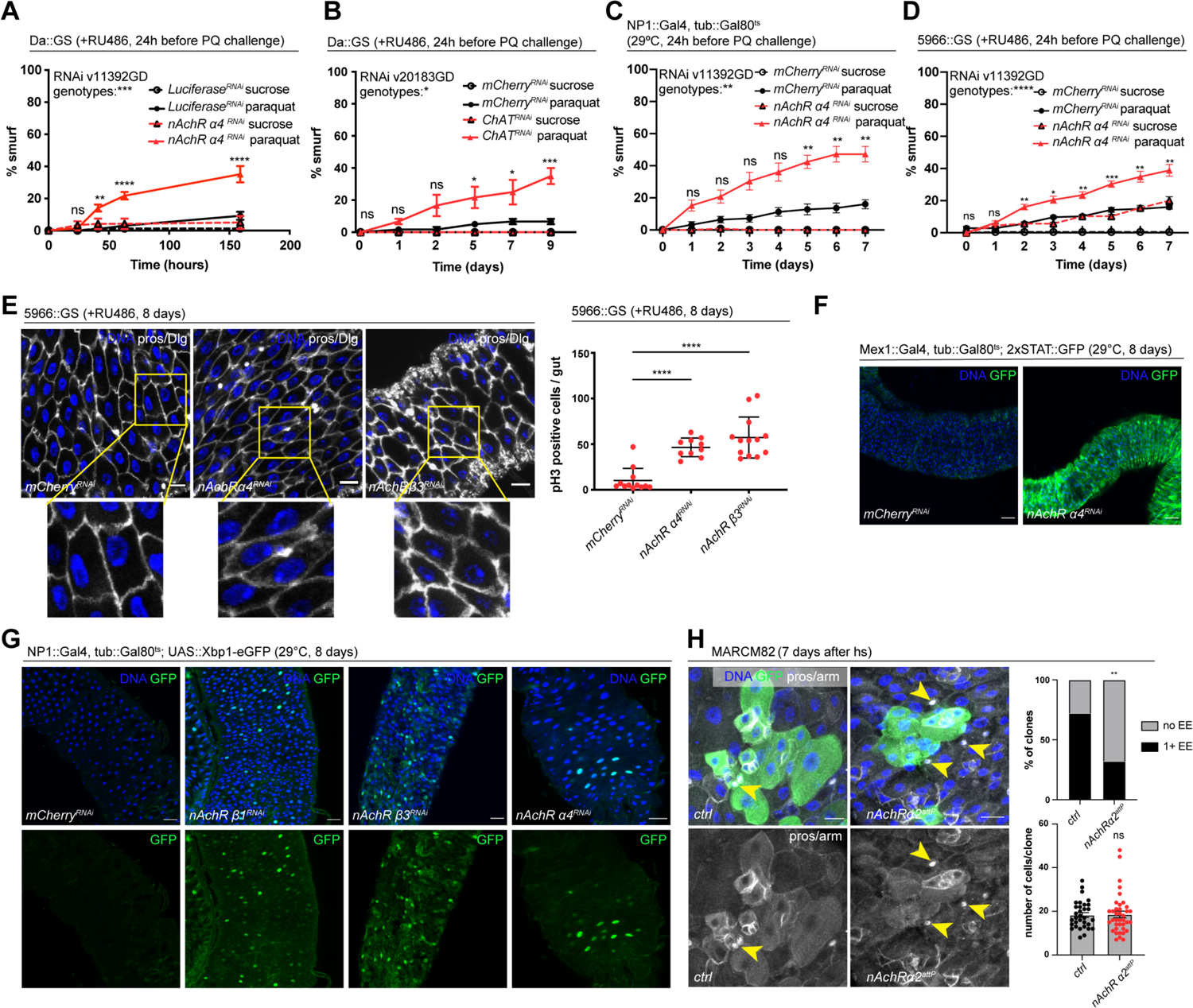
nAchR genes are required for barrier function in enterocytes (ECs) and enteroendocrine (EEs) cell differentiation. A) Barrier dysfunction assay after Luciferase (control) or nAchR *α*4 subunit depletion for 24h with ubiquitous driver Da::GS. nAchR *α*4: n=100 for Luciferase RNAi (control) on sucrose; n=125 animals for Luciferase RNAi on sucrose+paraquat; n=150 for nAchR *α*4 RNAi on sucrose; n=175 animals for nAchR *α*4 on sucrose+paraquat. Paraquat concentration 25mM. N=1. Two-way ANOVA followed by Šídák’s multiple comparisons test. B) Barrier dysfunction assay after mCherry (control) or ChAT depletion for 24h with ubiquitous driver Da::GS. n=75 animals per genotype and condition; N=3. Two-way ANOVA followed by Šídák’s multiple comparisons test. C) Barrier dysfunction assay after mCherry (control) or nAchR *α*4 depletion for 24h with enterocyte-specific driver NP1::Gal4, tub::Gal80^ts^ (NP1^ts^). n=125 animals per genotype and condition; N=3. Two-way ANOVA followed by Šídák’s multiple comparisons test. D) Barrier dysfunction assay after mCherry (control) or nAchR *α*4 depletion for 24h with enterocyte-specific driver 5966::GS. n=175 animals per genotype and condition; N=3. Two-way ANOVA followed by Šídák’s multiple comparisons test. E) Confocal immunofluorescence images examining epithelial organization and quantification of ISC mitoses in guts depleted of nAchR *β*3 and *α*4 subunits in ECs for 8 days. Septate junctions are stained with anti-Dlg antibody (white), DNA (blue) is labeled with Hoechst. Yellow boxed insets are shown enlarged in bottom row. Scale bars 10μm. Mitotically active ISCs are labeled with anti-pH3 antibody; n=12;10;13 guts for mCherry (control), nAchR *α*4 and nAchR *β*1, respectively. N=3. Ordinary one-way ANOVA followed by Dunnett’s multiple comparisons test. F) Confocal microscopy images of guts expressing a 2xSTAT::GFP reporter (green) depleted of mCherry (control) or nAchR *α*4 for 8 days in ECs with Mex::Gal4, tub::Gal80^ts^. n=10 guts per genotype. N=3. Scale bar 50μm. G) Confocal immunofluorescence image of posterior midguts expressing the UPR-reporter UAS-Xbp1-EGFP (green) after 8 days of nAchR subunit knockdown by RNAi. The EGFP tag is only in frame with the Xbp1(s) coding sequence after splicing using the unconventional splice site, which occurs under stress conditions. DNA (blue) is labeled with Hoechst. n=8 guts per genotype. N=3. Scale bar 25μm. H) Confocal immunofluorescence images of wildtype and nAchR *α*2 MARCM clones (green) 7 days after heat shock. Stem cells and enteroblasts are stained with anti-armadillo antibody (white); EEs are labeled with anti-prospero antibody (white, nuclear signal highlighted with yellow arrowheads) and DNA (blue) is labeled with Hoechst. Scale bars 15μm. Quantification of EE numbers within clones: n=32;38 clones for wildtype or nAchR *α*2, respectively from 3 pooled experiments. Fisher’s exact test. Quantification of cell numbers/clone: n=32;38 clones for wildtype or nAchR *α*2, respectively from 3 pooled experiments. Unpaired two-tailed t-test.

Acetylcholine (Ach) is the physiological ligand for nAchRs and is produced by ChAT, an enzyme that catalyzes the transfer of an acetyl group from coenzyme acetyl-CoA to choline (Taylor P., 1999). Modulation of total organismal Ach levels by RNAi-mediated silencing of ChAT under control of Da::GS also resulted in increased barrier dysfunction after paraquat exposure (Fig. 2B, Fig. S1B), further supporting the role of Ach/nAChR signaling in maintaining intestinal epithelial homeostasis.

To investigate a possible direct intestinal role for nAchR, and to identify the requirement for individual subunits, we used the drivers 5966::GS and NP1::Gal4 to separately deplete nAchR subunits. These drivers induce expression of UAS-linked transgenes in enteroblasts and enterocytes (Jiang et al., 2009; Zeng & Hou, 2015). While 5966::GS is inducible using RU486, we combined NP1::Gal4 with tub::G80ts (NP1ts) to allow for temperature-mediated induction (TARGET system (McGuire et al., 2004)) before subjecting the flies to paraquat. Knockdown of nAchR *α*4 or *β*3 with both drivers increased the numbers of smurf flies, indicating a defective epithelial barrier (Fig. 2C, D, Fig. S1C, D).

Knockdown of nAchR in ECs resulted in various hallmarks of epithelial stress. These include a mildly disorganized epithelial morphology (visualized by staining for the septate junction marker Dlg) and induction of intestinal stem cell (ISC) proliferation (Fig. 2E), presumably due to stress signals released by ECs (Biteau et al., 2011), as well as activation of JAK/STAT signaling (measured using the 10xSTAT::GFP reporter (Bach et al., 2007)) and ER stress signaling (measured using an Xbp1::GFP reporter (Sone et al., 2013) (Fig. 2F, G). Interestingly, Dlg staining did not indicate a strong disruption of EC junctions in the epithelium, suggesting that barrier dysfunction may be caused by a separate mechanism.

To further confirm and characterize the role for nAchR subunits in epithelial homeostasis, we generated MARCM mutant clones (Lee & Luo, 2001) lacking *nAChRα2* using the null allele nAChRα2^attP^ generated by CRISPR/Cas9 based homologous recombination resulting in the introduction of an attP site, 3xP3-RFP and a loxP site (Deng et al., 2019; Lu et al., 2022). Clone formation, growth, and cell composition also provide insight into a possible role of *nAChRα2* in ISC proliferation and differentiation. While *nAChRα2^attP^* clones grew to similar cell numbers as their control counterparts, they failed to produce normal numbers of EEs, as only 32% of *nAChRα2* clones contained at least 1 EE compared to 72% of clones in the control samples (Fig. 2H). In addition to a broader role for nAchR in maintaining barrier integrity, nAchR may thus also be required for the proper differentiation of EEs. Whether this effect on EE differentiation contributes to the barrier dysfunction remains unclear.

### Acetylcholine promotes barrier function

We sought to identify the source of Ach activating these receptors in the gut epithelium next. Because of its well understood role as a neurotransmitter, we initially focused on the innervation of the fly gut, which has been described previously (Cognigni et al., 2011). Expression of UAS::GFP under the control of ChAT::Gal4 confirmed that some of these neurons are indeed cholinergic (Fig. 3A, B’). Upon closer examination of the epithelium, we also noticed a small subset of prospero-positive EEs expressing GFP, predominantly located in the R4 and R5 regions of the midgut (Fig. 3B, C). In addition, labeling of guts expressing GFP under the control of neuron and EE-driver, prospero-Gal4, combined with tub::Gal80^ts^ (pros^ts^) with a ChAT antibody confirmed ChAT expression in a subset of EEs (Fig. 3D).

**Figure 3:**
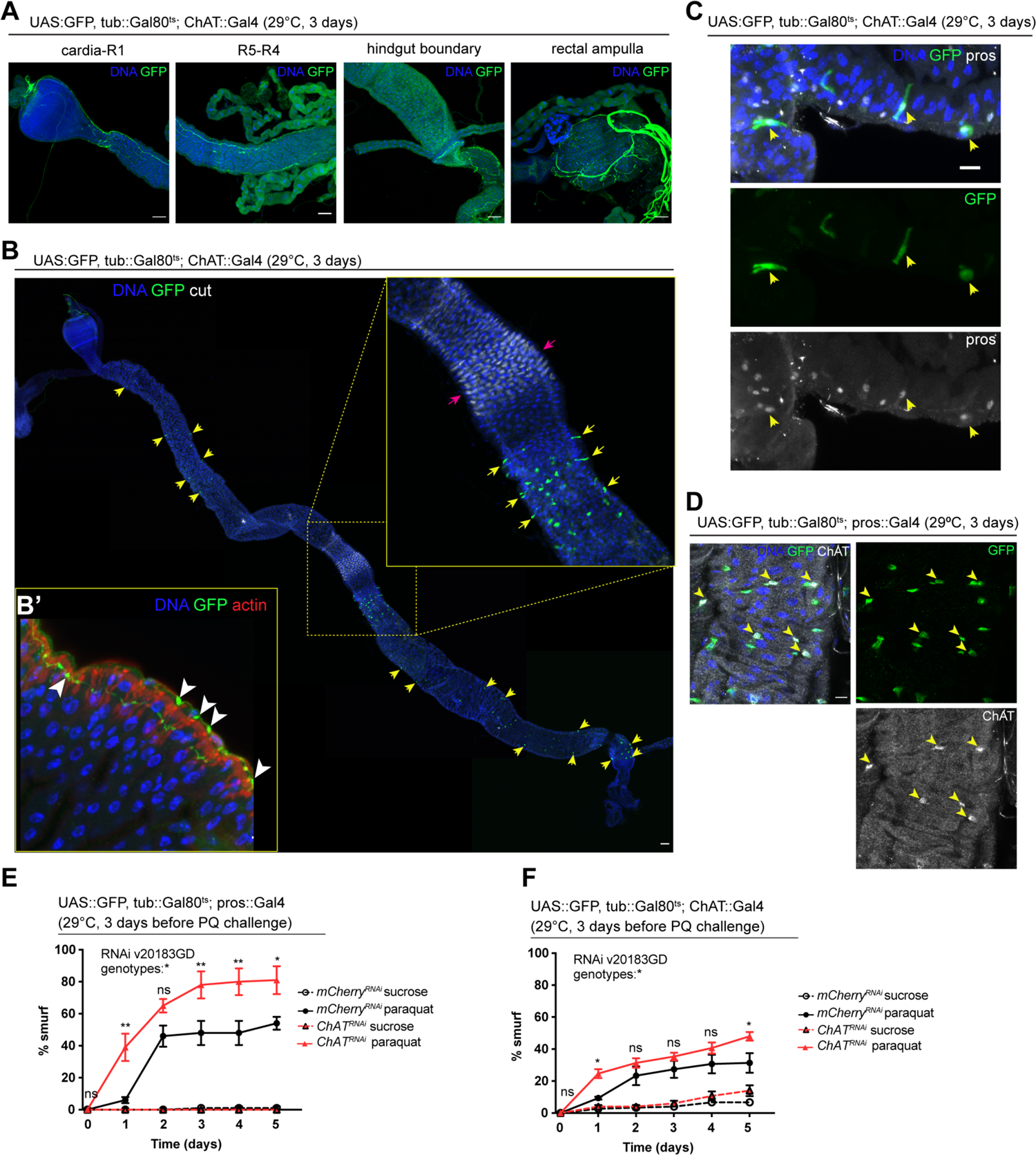
Acetylcholine produced in EEs and/or neurons promotes barrier function. A) Confocal immunofluorescence image of cholinergic innervation of different intestinal compartments. GFP (green) expression is driven by Mi{Trojan-GAL4.0}ChAT[MI04508-TG4.0] CG7715[MI04508-TG4.0-X] and detected in the anterior (cardia/R1) as well as posterior midgut (R4-R5), at the hindgut boundary and rectal ampulla. DNA (blue) is labeled with Hoechst. n=5 guts. N=3. Scale bars 50μm. B) Stitched confocal immunofluorescence images of a gut expressing GFP (green) under control of Mi{Trojan-GAL4.0}ChAT[MI04508-TG4.0] CG7715[MI04508-TG4.0-X], stained with anti-cut antibody (white). Yellow arrows indicate GFP-positive cells. Enlarged insert shows GFP-positive cells adjacent to the gastric region labeled with cut (pink arrows). DNA (blue) is labeled with Hoechst. n=5 guts. N=3. Scale bar 50μm B’) Confocal image of a gut expressing GFP (green) under control of Mi{Trojan-GAL4.0}ChAT[MI04508-TG4.0] CG7715[MI04508-TG4.0-X], stained with Phalloidin (red). Transverse section of the epithelium is shown revealing inter-epithelial axons from ChAT+ neurons. White arrowheads highlight axonal boutons. n=5 guts. N=3 C) Fluorescent immunohistochemistry image of posterior midgut expressing GFP (green) under the control of Mi{Trojan-GAL4.0}ChAT[MI04508-TG4.0] CG7715[MI04508-TG4.0-X], stained with anti-prospero antibody (white). Arrows indicate GFP-positive cells that also label for pros. DNA (blue) is labeled with Hoechst. n=8 guts. N=3. Scale bar 10μm. D) Confocal immunofluorescence image of ChAT antibody staining of the posterior midgut. EEs are expressing GFP (green) driven by pros::Gal4, yellow arrows indicate the overlap between ChAT staining (white) and pros-positive cells. DNA (blue) is labeled with Hoechst. n=8 guts. N=3. Scale bar 10μm. E) Barrier dysfunction assay after mCherry (control) or ChAT knockdown in EEs for 3 days with prospero-Gal4. n=100 animals per genotype and condition; N=3. Two-way ANOVA followed by Šídák’s multiple comparisons test. F) Barrier dysfunction assay after mCherry (control) or ChAT knockdown with Mi{Trojan-GAL4.0}ChAT[MI04508-TG4.0] CG7715[MI04508-TG4.0-X] for 3 days. n=120 animals per genotype and condition; N=3. Two-way ANOVA followed by Šídák’s multiple comparisons test. Data presented as mean ± SEM. ns, not significant, P > 0.05; *P ≤ 0.05; **P ≤ 0.01; ***P ≤ 0.001; ****P ≤ 0.0001. n: number of animals or midguts analyzed; N: number of independent experiments performed with similar results and a similar n

To address the role of Ach production in barrier integrity, we depleted ChAT with pros^ts^, as well as with ChAT::Gal4. Reduction of ChAT levels with both drivers rendered flies more susceptible to barrier dysfunction after paraquat exposure (Fig. 3E, F). Combined, these data support the notion that Ach signaling is critical to maintain barrier integrity and stress resilience in the intestinal epithelium of the fly. While cholinergic innervation is a likely source of the ligand in this response, local production of Ach by enteroendocrine cells may also play a role in maintaining homeostasis.

### Transcriptional changes after disruption of Ach signaling in the intestinal epithelium

As we observed barrier dysfunction without obvious deregulation of epithelial junctions after nAChR loss in ECs, we decided to profile changes in gene expression elicited in the gut by nAchR depletion. We performed RNAseq on whole guts depleted for nAchR *β*1 or *β*3 for 3 days under the control of NP1^ts^. PCA analysis suggested that the transcriptomes from intestines with nAchR knockdown were clearly distinct from transcriptomes of intestines with a control RNAi construct (mCherry RNAi; Fig. 4A). Overall, we observed 240 upregulated and 215 downregulated genes (Fig. 4B; FDR<= 0.1; log2(fold change) < −1 or > 1; 100% of samples have >= 1 reads), of which 171 were differentially expressed in both nAchR *β*1 and *β* knockdowns, supporting the idea that these subunits have partially overlapping functions (Fig. 4B, Fig. S2A, E). Synaptotagmin 4 (Syt4) was the most significantly downregulated gene in both knockdowns (Fig. 4D). GO term enrichment analysis revealed increased expression of glucosidases and hydrolases after nAchR knockdown (Fig. S2B, E), and downregulation of genes involved in immune responses such as lysozymes (Fig. 4C, Fig. S2C) and genes related to chitin binding and metabolism.

**Figure 4:**
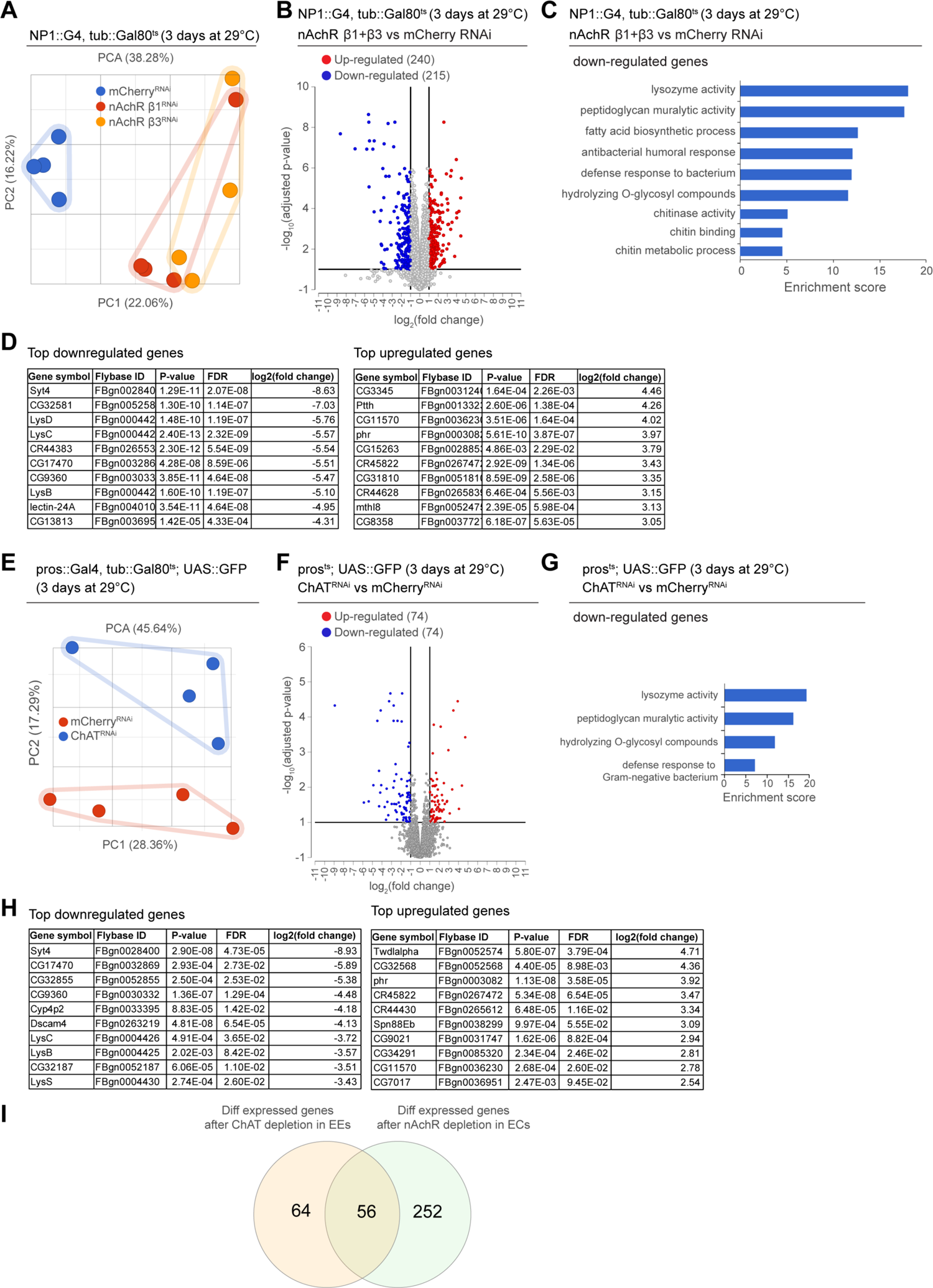
Transcriptional changes after disruption of Ach signaling in the intestinal epithelium. A) PCA plot of samples after 3 days of nAchR subunit depletion by RNAi in enterocytes with NP1^ts^. n=4 samples. N=1. B) Volcano plot showing significantly differentially regulated genes after short-term nAchR *β*1 or *β*3 knockdown in enterocytes. (FDR<= 0.1; log2(fold change) < −1 or > 1; 100% of samples have >= 1 reads) C) Gene set enrichment analysis of significantly downregulated genes after nAchR *β*1 and *β*3 knockdown in ECs. D) Top 10 most down- or upregulated genes after 3 days of nAchR subunit depletion by RNAi in enterocytes with NP1^ts^. E) PCA analysis after 3 days of ChAT depletion with RNAi in EEs under control of pros^ts^. n=4 samples. N=1. F) Volcano plot of significantly differently regulated genes after 3 days of ChAT knockdown in EEs. (FDR<= 0.1; log2(fold change) < −1 or > 1; 100% of samples have >= 1 reads) G) Gene set enrichment analysis of significantly downregulated genes after ChAT depletion in EEs. H) Top 10 most down- or upregulated genes after 3 days of ChAT knockdown by RNAi in EEs with pros^ts^. I) overlap between differentially regulated genes after 3 days knockdown of ChAT in EEs or nAchR *β*1 and *β*3 in ECs. n: number of samples included; N: number of independent experiments performed with similar results and a similar n.

In parallel, we analyzed the transcriptome of whole guts depleted of ChAT using pros^ts^ for 3 days. While the knockdown samples also separated clearly from the control, they displayed fewer differentially regulated genes than guts depleted of nAchR (Fig. 4E, F, Fig. S2F). However, Syt4 remained the most significantly downregulated gene (Fig. 4H) and enriched GO terms overlapped significantly with the previous experiment, especially with regards to immune responses as well as chitin metabolism (Fig. 4G, Fig. S2D). Direct comparison of differentially regulated genes revealed an overlap of 56 genes between guts depleted for nAchR in ECs and guts where ChAT was silenced using pros^ts^ (Fig. 4I).

### nAchR depletion disrupts PM integrity

The enrichment of chitin GO terms in our RNAseq experiments prompted us to examine the peritrophic matrix (PM). The PM is a protective structure lining the gut of many insects, consisting of crosslinked glycoproteins, proteoglycans and chitin (Erlandson et al., 2019; Hegedus et al., 2009; Hegedus et al., 2019). It surrounds the food bolus and forms a selectively permeable physical barrier preventing direct contact between abrasive food particles and bacteria with the epithelium, thus helping to compartmentalize digestive processes as well as protecting the animal from ingested toxins and pathogens (Erlandson et al., 2019; Hegedus et al., 2019). In flies, it was shown that the PM protects against pathogenic bacteria and their pore-forming toxins, such as *Pseudomonas entomophilia* and *Serratia marcescens* (Kuraishi et al., 2011). Dipteran insects such as *Drosophila* are thought to continuously produce a type II PM originating in the cardia at the anterior end of the midgut (Hegedus et al., 2019). There is evidence suggesting remodeling activity along the posterior midgut, as transcripts for PM components were found enriched in the R4 region of the midgut (Buchon et al., 2013). Moreover, intestinal IMD signaling as well as a subset of enteric neurons have been implicated in modulating the composition and permeability of the PM, however the underlying molecular mechanisms of PM remodeling remain poorly understood (Buchon, Broderick, Poidevin, et al., 2009; Kenmoku et al., 2016).

Transcript levels of two components of the PM, Crys and CG32302, were noticeably reduced in guts depleted of nAchR subunits (Fig. S3A). Earlier studies highlighted the importance of the PM in protecting the animal against lethal pathogenic bacterial infection with *Pseudomonas entomophila* (PE) (Kuraishi et al., 2011). Indeed, depletion of nAchR subunits in ECs significantly reduced survival after PE infection (Fig. 5A).

**Figure 5:**
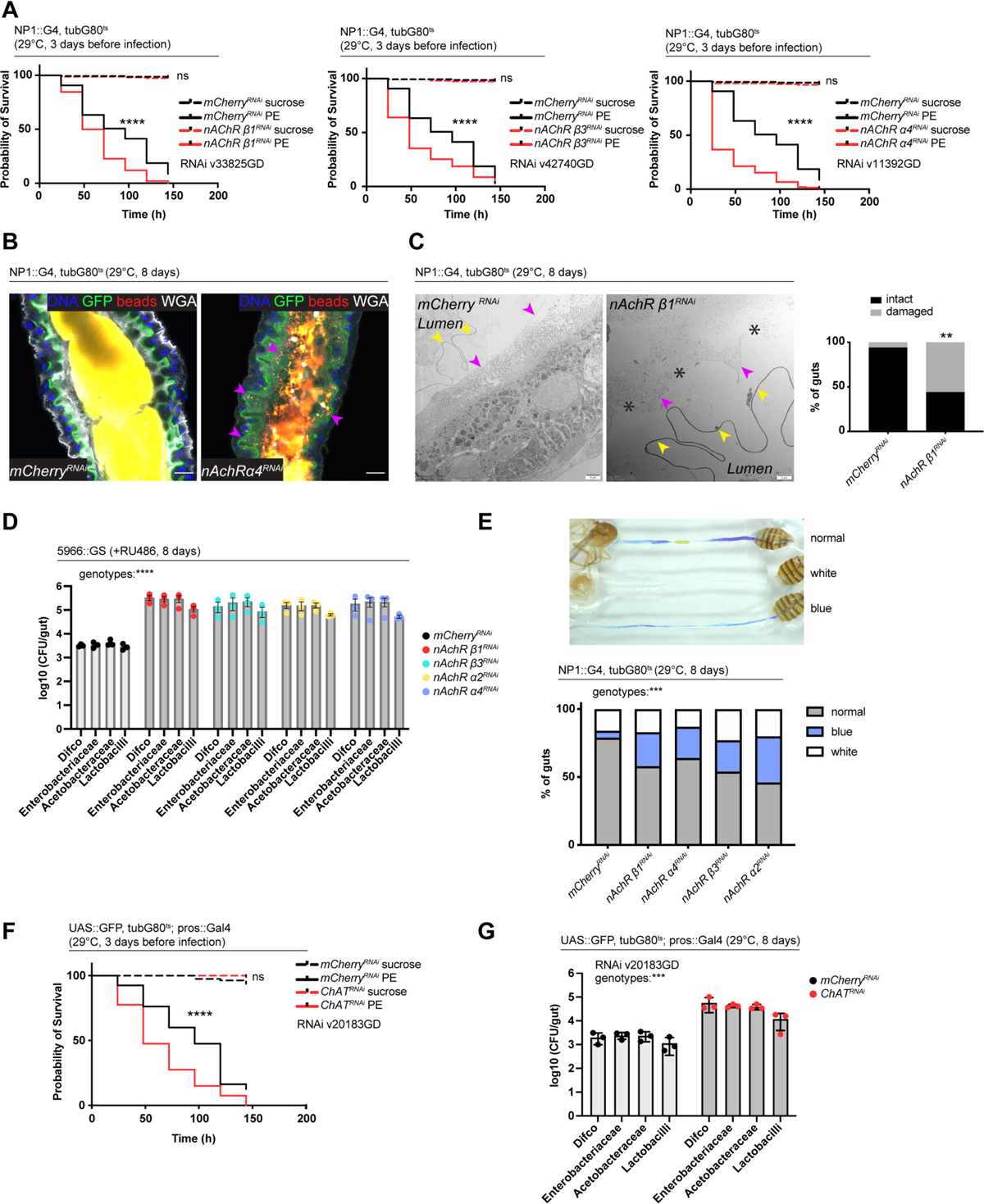
nAchR depletion disturbs PM integrity, causes dysbiosis and inflammation. A) Survival of animals depleted for mCherry (control) or nAchR *β*1, *β*3 or *α*4 for 3 days before *Pseudomonas entomophila* infection. n=150 animals per genotype and condition; N=3. Log Rank (Mantel-Cox) test. B) Confocal immunofluorescence image of posterior midguts depleted for either mCherry (control) or nAchR *α*4 for 8 days. Animals are expressing a GFP-brush border marker (green) and were fed red fluorescent beads to assess peritrophic matrix (PM) integrity (beads appear yellow/orange due to autofluorescence of beads in GFP channel). PM is labeled with WGA (white), DNA (blue) is labeled with Hoechst. Pink arrowheads highlight beads no longer contained by the PM sleeve. n=15 guts per genotype. N=3. Scale bar 20μm. C) Electron microscopy images and quantification of thin PM layer integrity. Thick (yellow arrows) and thin (pink arrows) PM layers are indicated. Asterisks highlight gaps in the thin layer after nAchR *β*1 depletion. n=16; 18 midguts. N=1. Fisher’s exact test. D) Colony forming units (CFU) of whole guts plated on selective growth media after 8 days of nAchR subunit depletion in ECs. 3 pooled independent experiments are shown. n=5 pooled animals per genotype and experiment. Two-way ANOVA. E) Gut compartmentalization and acidity after mCherry (control) or nAchR *β*1, *β*3 or *α*4 depletion for 8 days. Healthy flies fed with Bromphenol blue pH indicator display an acidic patch (yellow), while loss of gut compartmentalization leads to all blue or white guts. n=87 guts for mCherry (control), n=93 guts for nAchR *β*1, n=113 guts for nAchR *α*4, n=87 guts for nAchR *β*3 and n=90 guts for nAchR *α*2. 4 independent pooled experiments are shown. Chi square test. F) Survival after 3 days of mCherry (ctrl) or ChAT depletion in EEs followed by *Pseudomonas entomophila* infection. n=80 animals per genotype and condition; N=3. Log Rank (Mantel-Cox) test. G) Colony forming units (CFU) of whole guts plated on selective growth media after 8 days of ChAT depletion in EEs. 3 pooled independent experiments are shown. n=5 pooled animals per genotype and experiment. Two-way ANOVA. Data presented as mean ± SEM. ns, not significant, P > 0.05; *P ≤ 0.05; **P ≤ 0.01; ***P ≤ 0.001; ****P ≤ 0.0001. n: number of animals or midguts analyzed; N: number of independent experiments performed with similar results and a similar n.

Defects in the PM can be visualized with confocal light microscopy by feeding animals fluorescently labeled latex beads that are retained in the food bolus and stay separated from the epithelium if the PM sleeve is intact (Kenmoku et al., 2016). The surface of the bead-containing ingested food appeared relatively smooth in control animals. In contrast, silencing of nAchR *β*1 or *β*3 led to spiny protrusions of the fluorescent matter, indicating a damaged PM (Fig. S3B). We further modified this assay by crossing in the brush border marker A142::GFP (Buchon et al., 2013) and visualizing the PM with fluorescently labeled wheat germ agglutinin (WGA), a lectin that recognizes chitin (Carlini & Grossi-de-Sá, 2002), in addition to feeding the latex beads. Guts depleted of nAchR *α*4 displayed fluorescent signal scattered throughout the lumen and making contact with the brush border while the beads stayed confined to the PM sleeve and separated from the epithelium in control guts (Fig. 5B).

Electron microscopy has been successfully applied to detect subtle defects in PM morphology (Kuraishi et al., 2011). We therefore performed an ultrastructural analysis of the R4 compartment of guts depleted of nAchR *β*1. The PM was visible as a continuous folded ring in the lumen of control flies, consisting of electron-dense membranous material of roughly 100-200 nm thickness. Additionally, a second, much thinner (15-20nm) membranous ring-shaped layer was observed between the PM and the apical surface of the epithelial cells (Fig. 5C, Fig. S3C).

In the majority of the 18 examined nAchR *β*1 knockdown midguts the thick PM layer was not compromised (Fig. S3D). However, in 56% of samples the thin layer was clearly disrupted or missing altogether (Fig. 5C). Notably, none of the guts presented an intact thin layer in the absence of the thick layer. In all examined control and knockdown guts, the septate junctions connecting adjacent cells appeared normal, consistent with our Dlg staining, as well as the fact that no changes in junctional protein expression was observed in our RNAseq experiments.

### Defects in Ach signaling disturb gut compartmentalization and cause dysbiosis and JAK-STAT-mediated inflammation

Since the PM has been connected to regulation of the microbiome in mosquitos (Rodgers et al., 2017), we hypothesized that nAchR silencing also deregulates the microbial community inhabiting the fly gut. To test this assumption, we measured microbial load by plating pooled guts of control or nAchR knockdown animals on selective media supporting the growth of commensals such as Lactobacilli, Acetobacteriaceae or Enterobacteriaceae. The amount of CFUs after 8 days of nAchR *β*1, *β*3 or *α*4 silencing exceeded control levels significantly, indicating that these flies struggle to maintain appropriate commensal numbers (Fig. 5D). The fly midgut is functionally compartmentalized and contains a stomach-like region of acid-producing copper cells (Dubreuil, 2004). A previous study has highlighted the importance of gut compartmentalization in controlling microbiome abundance and distribution. As flies age, this spatial organization is progressively lost due to chronic JAK-STAT activation leading to metaplasia of copper cells in the acidic gastric region, ultimately resulting in dysbiosis and death of the animal (Li et al., 2016). Gut compartmentalization can be visualized by feeding flies the pH indicator Bromphenol blue, which labels the acidic copper cell region in yellow, while the rest of gut remains blue, indicative of a more basic pH (Li et al., 2016). Reduction of nAchR levels lead to an increase of disturbed acidity patterns, ranging from completely blue guts to samples with weak staining and white patches, which has been attributed to expansion of acid-producing commensals like Lactobacillus along the whole gut (Li et al., 2016) (Fig. 5E).

Dysbiosis can be a consequence of IMD pathway disruption, but at the same time triggers chronic IMD pathway activation (Buchon, Broderick, Chakrabarti, et al., 2009; Guo et al., 2014) Surprisingly, we did not observe an upregulation of antimicrobial peptides (AMP) transcripts classically associated with an IMD response (De Gregorio et al., 2002; Imler & Bulet, 2005). Furthermore, depletion of ChAT in EEs with pros^ts^ also caused increased susceptibility to PE infection (Fig. 5F) as well as dysbiosis (Fig. 5G). Conversely, overexpression of ChAT promoted survival after bacterial infection (Fig. S3E). Together these results suggest that Ach signaling is required to maintain a healthy microbiome and protect the animals against pathogenic infections.

### Syt4 is a transcriptional target of nAchR regulating PM function

One of the most significantly downregulated genes identified in our RNAseq data sets is Synaptotagmin 4, a vesicular Ca^2+^-binding protein promoting retrograde signaling at synapses (Yoshihara et al., 2005). RNAi-meditated silencing of Syt4 under control of NP1^ts^ reduced survival after challenge with PE (Fig. 6A, Fig. S4A) and caused PM defects visualized with the bead feeding assay (Fig. 6B). Moreover, Syt4 depletion increased commensal numbers (Fig. S4B), disrupted gut compartmentalization (Fig. S4C) as well as the morphology of the gastric region: Acid-producing copper cells usually form deep invaginations of the apical membrane (Dubreuil, 2004), giving rise to a gastric unit that can be visualized with anti-cut staining (Li et al., 2016). Syt4-depletion resulted in a disorganized morphology and a marked flattening of these units (Fig. S4D).

**Figure 6:**
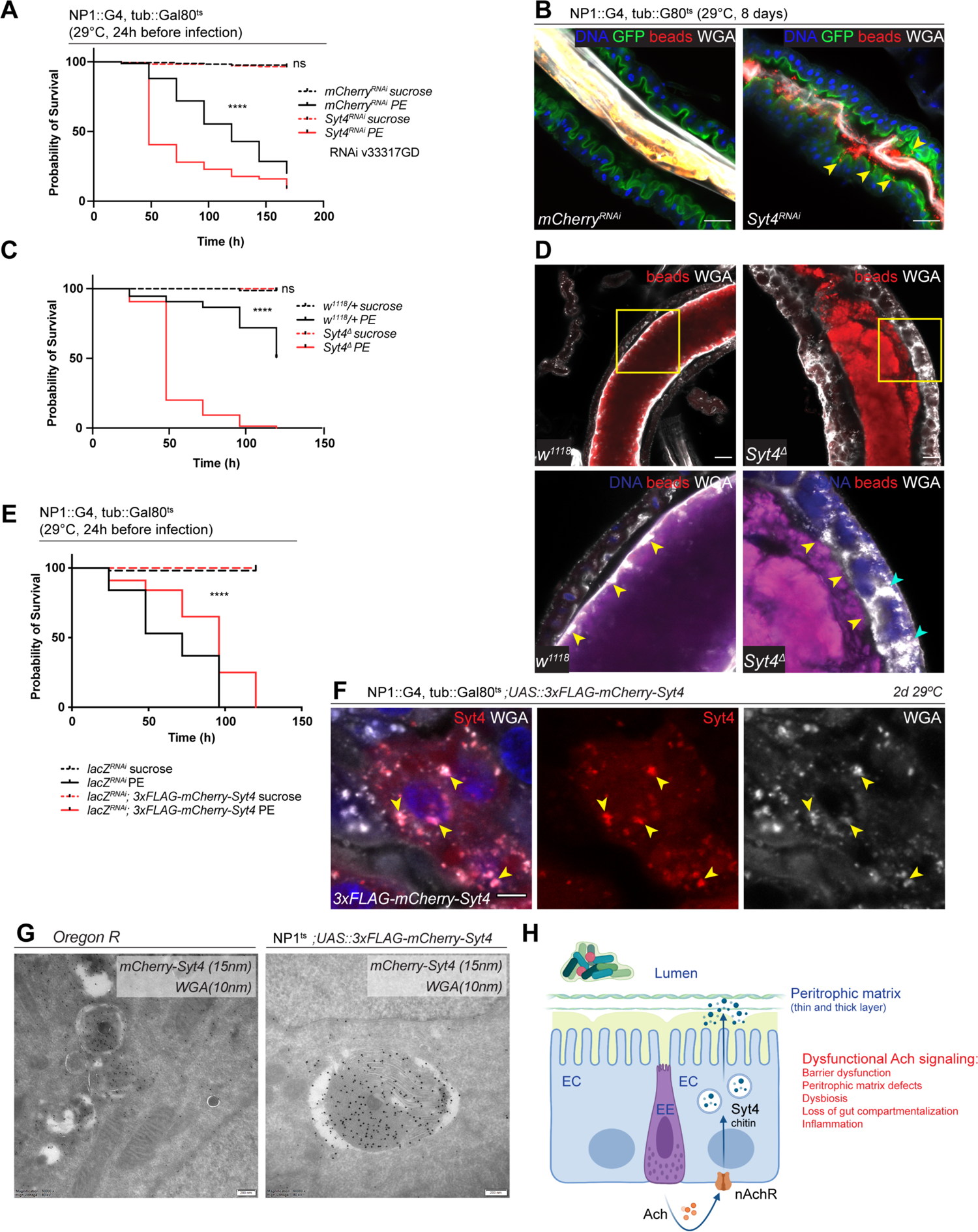
Syt4 knockdown affects PM integrity and phenocopies nAchR depletion. A) Survival after one day of mCherry (ctrl) or Syt4 depletion in ECs followed by *Pseudomonas entomophila* infection. n=175 animals per genotype and condition; N=3. Log Rank (Mantel-Cox) test. B) Confocal immunofluorescence image of posterior midguts depleted for either mCherry (control) or Syt4 for 8 days. Animals are expressing a GFP-brush border marker (green) and were fed red fluorescent beads to assess peritrophic matrix (PM) integrity (beads appear yellow/orange due to autofluorescence of beads in GFP channel). PM is labeled with WGA (white), DNA (blue) is labeled with Hoechst. Yellow arrowheads indicate beads that leaked out of the PM sleeve. n=10 guts per genotype. N=3. Scale bar 25μm. C) Survival of outcrossed w1118 (control) or Syt4^Δ^ CRISPR null mutant flies after *Pseudomonas entomophila* infection. n=75 animals per genotype and condition; N=3. Log Rank (Mantel-Cox) test. D) Confocal immunofluorescence image of posterior midguts of w1118 (control) or Syt4^Δ^ animals fed with red fluorescent beads to monitor PM integrity. PM is stained with WGA (white), DNA (blue) is labeled with Hoechst in bottom panels. Yellow insets are shown enlarged in bottom row. Yellow arrowheads indicate the presence (w1118) or absence of a clear PM boundary. Cyan arrowheads indicate accumulation of WGA signal within the epithelium. n=10 guts per genotype. N=3. Scale bar 25μm. E) Survival after overexpression of Luciferase-RNAi (control) or Luciferase-RNAi together with UAS-FLAG-mCherry-Syt4 for one day before *Pseudomonas entomophila* infection. n=100 animals per genotype and condition; N=3. Log Rank (Mantel-Cox) test. F) Confocal immunofluorescence image of posterior midguts overexpressing UAS-FLAG-mCherry-Syt4 (red) in enterocytes stained with WGA (white). DNA (blue) is labeled with Hoechst in bottom panels. Yellow arrowheads indicate overlap between Syt4-positive vesicles and WGA staining. n=8 guts. N=3. Scale bar 25μm. G) Immunogold electron microscopy image of posterior midgut of an Oregon R wildtype animal or an animal overexpressing UAS-FLAG-mCherry-Syt4 in enterocytes with NP1ts. WGA-biotin (10nm gold particles) is detected in multilamellar bodies carrying membranous and amorphous material. Syt4 (stained with anti-mCherry antibody, 15nm gold particles) colocalizes with these structures in animals expressing the UAS-FLAG-mCherry-Syt4, while Oregon R samples are devoid of anti-mCherry antibody labeling. n= 5. N=1. Scale bar 200nm. H) Model: Neuronal or EE-derived Ach maintains barrier function through Syt4-mediated secretion of PM components such as chitin from ECs. Disrupted Ach signaling leads to barrier dysfunction, peritrophic matrix defects, dysbiosis, as well as loss of gut compartmentalization and inflammation. Ach, Acetylcholine; nAchR, nicotinic acetylcholine receptor; EC, enterocyte; EE, enteroendocrine cell; Syt4, Synaptotagmin 4

We generated a new Syt4 null mutant with CRISPR/Cas9 technology to further substantiate these findings. While these mutant animals were homozygous viable, they displayed enhanced susceptibility to PE challenge (Fig. 6C) and a fragmented PM, often accompanied by enlarged WGA-positive structures within the epithelium (Fig. 6D).

To characterize the localization of Syt4 in the gut epithelium we utilized a 3xFLAG-mCherry-labeled protein trap line under UAS control (Singari et al., 2014). Overexpression of this construct with NP1^ts^ had a protective effect on animal survival after PE challenge (Fig. 6E) and yielded a vesicular staining pattern that overlapped with Golgi and lysosomal markers (Fig. S4E). Interestingly, we noticed a significant colocalization of Syt4-mCherry-positive structures and WGA staining in immunofluorescence experiments, suggesting that Syt4 vesicles contain chitin (Fig. 6F). Immunogold electron microscopy confirmed the colocalization of Syt4-mCherry and WGA in vesicular structures containing highly folded membrane swirls and amorphous cargo (Fig. 6G). These vesicles also stained positive for the late endosomal/lysosomal marker Lamp1 (Fig. S4F). Similar WGA/Lamp1 expressing structures were observed in wild-type (OreR) guts (Fig. 6G; Fig. S4F).

## Discussion

Our study identified 22 candidate COPD genes that disrupt barrier integrity in the fly and demonstrates the utility of the genetically accessible fly to screen candidate disease genes from human GWAS studies to provide mechanistic insight into their role in tissue homeostasis and pathophysiology. In particular, our results provide a role for nAchR signaling in maintaining intestinal barrier function. Since depletion of several nAchR subunits in enterocytes, or of ChAT in neurons and EEs leads to loss of barrier integrity and decreased survival after chemical or bacterial challenge, we propose that acetylcholine-mediated crosstalk between cholinergic neurons and/or EEs with ECs is critical to maintain intestinal epithelial homeostasis. This role of nAchR signaling is mediated by transcriptional regulation of Syt4 in ECs, which in turn maintains secretion of chitin vesicles to maintain the peritrophic matrix (Fig. 6H).

A role for acetylcholine signaling in human barrier epithelia is supported by previous studies: muscarinic acetylcholine receptors have been successfully targeted in the clinic to relieve bronchoconstriction and mucus hypersecretion in COPD and asthma (Calzetta et al., 2021), although nicotinic AchRs remain more elusive from a therapeutic perspective (Hollenhorst & Krasteva-Christ, 2021). While Ach is a classic neurotransmitter, a growing body of work has uncovered an important role of Ach beyond the context of the nervous system: various non-neuronal cell types express the machinery for Ach synthesis and secretion, ranging from diverse immune cells to epithelial cells, such as brush/tuft cells (Kummer & Krasteva-Christ, 2014; Wessler & Kirkpatrick, 2008). Airway tuft cells have been implicated in orchestrating type 2 inflammatory responses (O’Leary et al., 2019; Sell et al., 2021) and mucociliary clearance (Perniss et al., 2020), whereas their intestinal counterparts participate in defense against helminths and protists and limit biliary inflammation (O’Leary et al., 2022; O’Leary et al., 2019). With a wide range of cell types able to produce or sense Ach, non-neuronal Ach serves as a versatile signaling molecule eliciting complex intercellular crosstalk with diverse outcomes; depending on the context, Ach may promote inflammation or conversely exert anti-inflammatory functions (Hollenhorst & Krasteva-Christ, 2021; Kummer & Krasteva-Christ, 2014; Sell et al., 2021). Accordingly, it was recently reported that expression of a COPD risk allele of CHRNA5 in epithelial cells leads to airway remodeling *in vivo*, increased proliferation and production of pro-inflammatory cytokines through decreased calcium entry and increased adenylyl-cyclase activity (Routhier et al., 2021).

Previous work has further shown a downregulation of junctional proteins such as ZO-1 and p120 after depletion of CHRNA5 in A549 cells lung cancer cells (Krais et al., 2011). While we observed a slightly disorganized pattern of junctional markers such as Dlg after nAchR subunit knockdown in the fly intestinal epithelium, junctional architecture appeared normal when analyzed by electron microscopy. Furthermore, transcriptome analysis revealed little to no changes in the expression of proteins involved in polarity or cellular junction formation, suggesting that nAchR signaling regulates barrier function through other mechanisms. A role for the Syt4-mediated secretion of PM protein components and chitin in maintaining barrier integrity is supported by the observation that mCherry-tagged Syt4 partially overlaps with chitin-binding wheat germ agglutinin staining.

While the *Drosophila* PM is thought to be produced mostly in the anterior most portion of the gut (Hegedus et al., 2019), the existence of WGA-positive vesicles throughout the entire midgut suggests continuous remodeling along the length of the tissue. This finding is consistent with the previously reported expression of PM-related transcripts in the R4 compartment of the midgut (Buchon et al., 2013). PM integrity can be modulated by enteric neurons, although a role for cholinergic signaling was not tested in this context (Kenmoku et al., 2016).

Our study highlights the evolutionary conservation of mechanisms maintaining epithelial barrier function. The PM is functionally analogous to mucus and surfactant layers in mammalian airways, and it remains to be explored whether COPD risk alleles in nAchR subunits also cause a dysfunction in the secretion of such barrier components. The elevated inflammation and airway remodeling in mice expressing the CHRNA5 risk allele suggest that such a mechanism may be conserved as well (Routhier et al., 2021). It is critical to note that the epithelial dysfunction observed in these animals, as well as part of the association of COPD risk with specific CHRNA loci, emerge independently of cigarette smoke (Parker et al., 2019; Routhier et al., 2021; Siedlinski et al., 2013), indicating that nAchR signaling is critical to maintain homeostasis not only in the context of oxidative stress, but under homeostatic conditions. Supporting this view, our data show that knockdown of nAchR subunits in fly ECs also results in epithelial stress signaling in the absence of Paraquat exposure.

These consistencies further validate the approach of prioritizing candidate genes associated with COPD risk loci using the *Drosophila* intestine as a model system. Characterization of the epithelial role of other identified candidate orthologues from our screen will likely provide further insight into the biology and pathophysiology of barrier dysfunction and epithelial homeostasis. Such studies will be critical for target identification and validation for therapeutic intervention in COPD.

## Material and methods

### *Drosophila* stocks and husbandry

Flies were raised and kept on standard fly food prepared according to the following recipe: 1 l distilled water, 13.8 g agar, 22 g molasses, 75 g malt extract, 18 g inactivated dry yeast, 80 g corn flour, 10 g soy flour, 6.26 ml propionic acid, 2 g methyl 4-hydroxybenzoate in 7.2 ml of ethanol. Flies were reared at 25°C with 65% humidity on a 12 h light/dark cycle. All animals used in this study were mated females matured for 4-6 days.

The TARGET system was used to conditionally express UAS-linked transgenes in specific cell populations in combination with indicated Gal drivers (McGuire et al., 2004). Crosses containing tub::Gal80^ts^ were reared at 18°C to avoid premature gene expression. Transgene expression was induced by shifting the flies to 29°C for 1-8 days, as indicated in the figure legends.

For experiments using a GeneSwitch driver, flies were reared on normal food before being shifted to food containing 200mM Mifepristone (RU486); for barrier function experiments (smurf assay). FD&C blue dye (Neta Scientific, SPCM-FD110-04) was added to the food at final concentration of 2.5% (w/v).

Formation of MARCM clones was induced with heat shock for 1h at 37°C and clones were analyzed after 7 days at 25°C.

RNAi lines used in the barrier dysfunction screen are listed in Suppl Fig. 1B.

The following additional lines were obtained from the Bloomington *Drosophila* Stock Center: w^1118^, Oregon-R, UAS-mCherry^RNAi^ (35785), UAS-ChAT^RNAi^ (60028, 25856), FRT82B (2051), w*; TI{TI}nAChRα2attP/TM6B (84540), w[*]; Mi{Trojan-GAL4.0}ChAT[MI04508-TG4.0] CG7715[MI04508-TG4.0-X]/TM6B (60317) The following additional lines were obtained from the Vienna *Drosophila* Stock Center: UAS-nAchRβ1^RNAi^ (33825, pruned), UAS-nAchRβ3^RNAi^ (42742, pruned), UAS-ChAT^RNAi^(20183, 330291). The following lines were gifts: 5966::GeneSwitch (B. Ohlstein), 2xSTAT::GFP (E. Bach), NP1::Gal4 (D. Ferrandon), A142:GFP (N. Buchon), Mex1::Gal4;tub::G80ts (L. O’Brien), MARCM82 (hsFlp; tub::Gal4, UAS::GFP; FRT82, tub::Gal80, Norbert Perrimon), ProsV1::Gal4 (PMID: 11486507), Da::GeneSwitch (PMID: 19486910), UAS::Xbp1-eGFP (H. D. Ryoo)

### Generation of UAS-ChAT

DNA encoding the sequence of Choline O-acetyltransferase (Uniprot identifier P07668, amino acid residues 1-721), was synthesized and subcloned into pUASTattB under the control of the hsp70 promoter. Transgenic lines were established by WellGenetics, Taiwan. In brief, pUASTattB plasmid containing the ChAT sequence was microinjected into embryos of y[1] M{vas-int.Dm}ZH-2A w[*]; P{y[+t7.7]=CaryP}attP40 or y[1] M{vas-int.Dm}ZH-2A w[*]; P{y[+t7.7]=CaryP}attP2. Transgenic F1 flies were screened for the selection marker white+ (orange colored eyes).

### Syt4 CRISPR mutant

CRISPR mediated mutagenesis was performed by WellGenetics, Inc. (Taiwan) using modified methods of Kondo and Ueda (Kondo & Ueda, 2013). In brief, the upstream gRNA sequences TTTCCACTCGATGTTCCTGG[CGG] and downstream gRNA sequences CGCAGGCGCCCCTTAATGAG[GGG] were cloned into U6 promoter plasmids separately. Cassette 3xP3 RFP, which contains a floxed 3xP3 RFP and two homology arms, were cloned into pUC57 Kan as donor template for repair. Syt4/CG10047-targeting gRNAs and hs Cas9 were supplied in DNA plasmids, together with donor plasmid for microinjection into embryos of control strain w[1118]. F1 flies carrying the selection marker 3xP3 RFP were further validated by genomic PCR and sequencing. This CRISPR editing generates a 2,603 bp deletion allele of Syt4, deleting the entire CDS and replacing it with a 3xP3 RFP cassette.

### Gene assignment to COPD genome-wide association study (GWAS) loci

Publicly available summary statistics for the discovery stage of the COPD GWAS reported by Hobbs *et al.,* 2017 were obtained from dbGaP (accession: phs000179). Forty-eight candidate genes were assigned to the 22 loci reported in Hobbs *et al.,* 2017 based upon expression quantitative trait loci (eQTL), coding variation level support or physical distance if a gene could not be assigned via the former criteria. First, candidate genes were assigned to loci if the index variant was an eQTL in any tissue for any gene within 250 kilobases of the variant in GTEx (Battle et al., 2017) (V6p). We further applied colocalization (via the coloc package in R) (Giambartolomei et al., 2014) to estimate the probability the eQTL and COPD risk association signal share a casual variant. Of the 40 genes with eQTL support, 24 had a colocalization probability > 0.6. Candidate genes were also assigned to loci if the index variant was in linkage disequilibrium (LD) (r^2^>0.6) with coding variants for the gene. LD was estimated using individuals of European ancestry from 1000 Genomes (Auton et al., 2015). Eight candidate genes were assigned to five loci, six of which overlapped genes with eQTL level support.

Since we first obtained this candidate gene list, a larger COPD risk GWAS was published (Sakornsakolpat et al., 2019) that made use of not only lung eQTL and coding variant data, but also epigenetic and gene-set similarity approaches to assign candidate genes to COPD risk loci (see Supplementary Table 7 in Sakornsakolpat *et al*., 2019). We found our assigned candidate genes overlapped with candidate genes from this newer study at 13/22 loci reported in the Hobbs et al., 2017 study, including CHRNA3/5. Overall 20/48 candidate genes were also listed as candidate genes in the Sakornsakolpat et al. study.

### Barrier Dysfunction Screen

For the barrier dysfunction assay, males from candidate RNAi lines were crossed at a 1:1 ratio with virgin *Daughterless* (*Da*)::GeneSwitch driver line females in Bloomington-modified food (standard medium) bottles. Crosses were raised at 25°C and brooded every 2-3 days. Progeny were collected and females were sorted after mating for 2-3 days (discarding males). This yielded about 200 females per genotype depending on the RNAi line. Sorted females were aged in standard medium at 25°C for 10-12 days. Aged females (∼25 per vial; exact number recorded per vial for assay read-out) were then exposed to standard medium prepared with 200mM Mifepristone (RU486) from Sigma Aldrich (cat# 856177) and 2.5% w/v FD&C Blue Dye no 1 from Spectrum (cat# FD110) for 24 hours at 25°C or 29°C. Prior to paraquat exposure, flies were dry starved for 2-3 hours at the experimental temperature. 25mM paraquat solution (5% sucrose and 2.5% w/v FD&C Blue Dye in sterile water) or mock solution (sucrose and blue dye only) was freshly prepared for each experiment.. Starved flies were placed in empty vials with a Whatman filter paper (VWR, 89013-946) on top of a foam biopsy pad (Neta Sciences, BPLS-6110) and 1.25mL of paraquat or mock solution for 16 hours at 25°C or 29°C and then shifted back to medium with 200mM Mifepristone (RU486) and 2.5% w/v FD&C Blue Dye no 1. Entirely Blue (Smurf) flies were counted starting post-16-hour exposure. Smurf flies were counted daily or every-other day. About 8-12 candidate RNAi lines were tested in sets with Luciferase RNAi always included as a control.

The average proportion of smurf flies across technical replicates per time point were calculated and graphed. The natural log (LN) ratio was calculated for each candidate RNAi by dividing the candidate RNAi proportion average from the final time point by the luciferase RNAi proportion average for the same time point (=LN(Candidate RNAi/Luciferase RNAi)). Candidate RNAi results were ranked by establishing a scale with arbitrary LN ratio ranges to define: strong enhancers (≥ 1), enhancers (≥ 0.43 to ≤ 0.99), weak enhancers (≥ 0.20 to ≤ 0.42), no effect (≥ −0.30 to ≤ 0.19), weak suppressors (≥ −0.70 to ≤ −0.31), suppressors (≥ −1.10 to ≤ −0.71), and strong suppressors (≤ −1.11).

### Paraquat feeding

20-25 flies per vial were kept on food containing FD&C blue for 1-3 days and dry starved in empty vials for 2-3h prior to Paraquat exposure. Methyl viologen dichloride hydrate (Paraquat, 856177, Sigma Aldrich) solution was prepared freshly for each experiment in 5% (w/v) sucrose in water with 2% (w/v) FD&C blue. Paraquat concentration was 12.5mM unless indicated otherwise. Starved flies were transferred to vials containing 600μl of Paraquat solution or 5% sucrose (mock treatment) as well as a circular Whatman filter paper (VWR, 89013-946) on top of a foam biopsy pad (Neta Sciences, BPLS-6110). Flies were treated for 16h overnight and then transferred back to fly food with FD&C blue dye. The number of smurf flies was recorded 24h after the start of the Paraquat challenge and subsequently monitored over the course of 7-10 days.

### Pseudomonas entomophila infection

*Pseudomonas entomophila* (PE, gift from B. Lemaitre), was cultured in LB medium at 29 °C overnight for 16-18h (15ml/sample to be infected). Bacteria were centrifuged at 4000g for 10 min at RT and resuspended in 5% sucrose (OD_600_=100). 500μl of concentrated bacterial suspension or 5% (w/v) sucrose solution (mock treatment) was added to empty fly vials containing a circular Whatman filter paper (VWR, 89013-946) on top of a foam biopsy pad (Neta Sciences, BPLS-6110). 20-25 flies per vial were starved in empty vials for 2–3 h before infection. Survival was monitored over the course of 7-10 days and 100μl of sucrose solution was added daily.

### Gut compartmentalization

Gut compartmentalization was assessed as described in Li et al., 2016: 100μl of 2% w/v Bromphenol blue solution (Sigma Aldrich, B5525) was dispensed in a food vial, the surface was broken up with a pipette tip to allow full absorption of the dye before flies were transferred onto food. Flies were fed overnight and guts were dissected in small groups and immediately scored visually under a stereomicroscope to avoid prolonged exposure to CO_2_.

### Immunofluorescence microscopy

Guts from adult female flies were dissected in PBS, fixed for 45mins at room temperature (RT) in fixative solution (4% formaldehyde, 100 mM glutamic acid, 25 mM KCl, 20 mM MgSO_4_, 4 mM Na_2_HPO_4_, 1 mM MgCl_2_, pH 7.5), washed twice in wash buffer (1× PBS, 0.5% bovine serum albumin and 0.1% Triton X-100, 0.005% NaN_3_) for 30 min at RT. Primary and secondary antibodies were diluted in wash buffer. Samples were incubated overnight at 4°C with primary antibody, washed twice for 30min with wash buffer before incubating 4-6h at RT with secondary antibody. Hoechst33342 (Invitrogen, H3570, 1:1000) or wheat germ agglutinin-AlexaFluor647 (Invitrogen, W32466, 1:500) were added to the secondary antibody cocktail to visualize DNA or the peritrophic matrix (PM) respectively. Samples were washed again twice for 30mins before mounting in Prolong Glass antifade mounting media (Invitrogen, P36982).

To assess the integrity of the PM, flies were dry starved for 2h and then fed Fluoresbrite microspheres (Polysciences, 17149 (0.05μm, green) or 195075 (0.5μm, red), diluted 1:50 in 5% sucrose solution on Whatman filter paper for 16h overnight. Guts were dissected, fixed and processed for immunofluorescence microscopy analysis as described above.

For lysotracker staining, freshly dissected guts were incubated for 5mins in 1× PBS with Lysotracker Deep Red (Invitrogen, L12492, 1:500) before fixation. Samples were washed twice for 10mins before and after 1h incubation with Hoechst, mounted and analyzed within one day.

Primary antibodies used in this study: chicken anti-GFP (Abcam, ab13970, 1:1000), mouse anti-armadillo (DSHB, N2 7A1, 1:100), mouse anti-Delta (DSHB, C594.9B, 1:50), mouse anti-Dlg (DSHB, 4F3 anti-discs large, 1:20), mouse anti-ChAT (DSHB, ChAT4B1, 1:100), rabbit anti-phospho Histone H3 (Millipore, 06-570, 1:2000), mouse anti-prospero (DSHB, MR1A, 1:250), mouse anti-cut (DSHB, 2B10, 1:100), mouse anti-Golgin84 (DSHB, Golgin 84 12-1, neat) Secondary antibodies were from Jackson ImmunoResearch Laboratories and diluted 1:1000. donkey anti-mouse Cy3 (Jackson ImmunoResearch Laboratories, 715-165-150, 1:1000), donkey anti-mouse Alexa-647 (Invitrogen, A31571, 1:1000), donkey anti-chicken Alexa-488 (Jackson ImmunoResearch Laboratories, 703-545-155, 1:1000), donkey anti-rabbit Cy3 (Jackson ImmunoResearch Laboratories, 711-165-152, 1:1000), donkey anti-rabbit Alexa 647 (Jackson ImmunoResearch Laboratories, 711-605-152, 1:1000) All Images were taken on a Leica SP8 confocal microscope and processed using FIJI(Schindelin et al., 2012) and Adobe Illustrator.

### CFU counting

Commensal bacteria were cultured as described in (Guo et al., 2014):. In brief, intact flies were sanitized in 70% ethanol for 1min and rinsed 3x in sterile PBS. 5 guts per sample were dissected and homogenized in 250μl sterile PBS. Serial dilutions were plated on selective media, plates were incubated for 48-72h at 29°C and colonies counted.

Selective plates were prepared according to the following recipes: *Acetobacteriaceae*: 25 g/l D-mannitol, 5 g/l yeast extract, 3 g/l peptone, and 15 g/l agar. *Enterobacteriaceae*: 10 g/l Tryptone, 1.5 g/l yeast extract, 10 g/l glucose, 5 g/l sodium chloride, 12 g/l agar. *Lactobacilli* MRS agar: 70 g/l BD Difco Lactobacilli MRS agar. *Nutrient Rich Broth*: 23 g/l BD Difco Nutrient agar. All media were autoclaved at 121 degrees Celsius for 15 min.

### Electron microscopy (EM)

For the localization of Syt4, flies were allowed to express UAS-3xFLAG-mCherry-Syt4 in enterocytes under control of NP1ts for 2 days before dissection in PBS at room temperature. The dissected gut was cut with a sharp blade at the R3-R4 border and the R4-hindgut segment was immediately immersed in either one of 2 fixative solutions, to obtain samples for immuno-EM and for conventional EM in Epon-embedded material.

For immuno-EM (Slot & Geuze, 2007), the fixation was performed with 4% paraformaldehyde (PFA), 0.1% glutaraldehyde (GA) in PHEM buffer (60mM PIPES, 25 mM HEPES, 2 mM MgCl_2_, 10 mM EGTA), pH 6.9, for 1h at room temperature.

Subsequently, fixation was continued in 0.6 % PFA in PHEM buffer at 4 °C for several days. The samples were then rinsed in PBS, blocked with 0.15 % glycine in PBS, and gradually embedded in gelatin, from 2% (30 min) over 6% (30 min) to 12 % gelatin. Small blocks of solidified gelatin each containing 1 gut segment were cryoprotected overnight with 2.3 M sucrose. They were mounted on aluminum pins in a direction to expose the transversal cut at the R4 segment for cryo-ultramicrotomy and frozen in liquid N2. Syt4 was localized on ultrathin cryosections with polyclonal rabbit anti-RFP antibody (600-401-379, Rockland). Chitin was localized with biotinylated wheat germ agglutinin (B-1025-5, Vector laboratories) followed by polyclonal rabbit anti-biotin antibody (100-4198, Rockland). *Drosophila*-specific rabbit anti-Lamp1 antibody was a gift from Andreas Jenny (Chaudhry et al., 2022). Antibodies were detected with protein A-conjugated with 15 or 10 nm gold particles in a JEOL JEM-1011 electron microscope.

For conventional EM the fixation was performed in 2.5 % GA in 0.1 M Sorensen’s phosphate buffer (PB), pH 7.4, for 4 h at room temperature, then overnight at 4 °C. Subsequently, fixation was continued in 0.6 % PFA in 0.1 M PB at 4 °C for several days. After rinsing in 0.1 M PB, the guts were postfixed with 1 % OsO_4_ and 1.5 % K_3_[Fe(CN)_6_] in 0.07 M PB, stained en bloc in aqueous 0.5 % uranyl acetate, dehydrated in acetone and embedded in Epon. Transverse sections of the R4 gut segments were stained with uranyl acetate and lead citrate and examined in a JEOL JEM-1011 electron microscope.

### Bulk RNAseq

For bulk RNAseq analysis 4 independent biological replicates per sample consisting of 20-25 guts each were dissected and collected on dry ice. RNA was extracted using the Qiagen RNeasy Mini kit. cDNA was generated from 2 ng of RNA using the Smart-Sq V4 Ultra Low Input RNA Kit (Takara, 634894). Next, 150 pg of cDNA was used to produce sequencing libraries with the Nextera XT DNA Sample Preparation Kit (Illumina, FC-131-1024). Libraries were then sequenced for 50 single read cycles and 30 million reads per sample on Illumina NovaSeq 6000. Reads were aligned to the *Drosophila* genome, version BDGP6, using the GSNAP aligner as part of the HTSeqGenie R package (version 4.2). Reads that uniquely aligned within exonic boundaries of genes were used to derive expression estimates. nRPKM values, in which total library sizes were normalized using the median ratio method as previously described (Anders & Huber, 2010), were generated for each gene. Partek Flow was used to perform differential gene expression and PCA analysis, gene ontology term enrichment as well as creation of illustrative graphs.

### Statistical analyses

Generation of graphs and statistical analyses were performed with Graphpad Prism 9. Experiments with two conditions were compared with a two-tailed parametric Student’s T-test or Fischer’s exact test, as appropriate. Experiments with multiple conditions were evaluated either by ordinary one-way ANOVA followed by Dunnett’s post-hoc test to compare a control group with experimental conditions or a Chi square test for categorical data. Barrier dysfunction curves were analyzed with 2-way ANOVA followed by Sidak’s multiple comparisons test. Survival curves were compared with the Mantel-Cox method.

No statistical methods were used to predetermine sample sizes; sample sizes were determined based on variation observed in pilot experiments and previously published literature. Exact numbers are listed in figure legends. All animals were randomly allocated to treatment groups. The experimenter was blinded for image analysis and other quantifications. The number of repeats for each experiment is listed in figure legends, all attempts at replication were successful. The initial screen as well as electron microscopy and RNAseq experiments were performed once for data gathering and hypothesis generation; the data was later validated by other methods. No data points were excluded from analyses.

### Illustrative model

Illustrative model summarizing results was created with BioRender.com

### Data availability

All data generated and analyzed are included in the manuscript, figures and figure supplements. All sequencing data generated in this study will been deposited in GEO under accession code XX and XX.

### Illustrative model

Illustrative model summarizing results was created with BioRender.com

### Material availability

Fly lines generated in this study (UAS-ChAT and Syt4 CRISPR mutant) are available upon request.

### Adherence to community standards

This study and manuscript were prepared in accordance with ARRIVE and ICMJE guidelines.

## Acknowledgements

We thank Dr Andreas Jenny (Albert Einstein College of Medicine) for the *Drosophila*-specific rabbit anti-Lamp1 antibody. We thank Dr Aniek Janssen and Dr Lucie van Leeuwen (UMC Utrecht) for the Oregon R flies for EM. The microscopy infrastructure in this work is subsidized by the Roadmap for Large-Scale Research Infrastructure (NEMI) of Netherlands Organisation for Scientific Research (grant number 184.034.014 to J.K.).

## Summary of Supplemental Materials

### Supplemental Materials

**Table S1 (Related to Figure 1)** A) List of candidate genes for genetic variants (human) associated with COPD in Hobbs et al., 2017. Genes highlighted in blue had a clear *Drosophila* ortholog and were included in the screen. Abbreviations used: SNP, Single nucleotide polymorphism; CHR, chromosome; BP, base pair (GRCh37); eqtl, expression quantitative trait loci; Risk allele, allele associated with increased COPD risk; Alt allele, alternative allele; OR stage1, Odds-ratio of risk allele in stage 1 of Hobbs *et al*., 2017; P.stage1, P-value in stage 1 of Hobbs et al; P.meta, meta-analysis P-value in Hobbs et al; Evidence.Sakornsakolpat, evidence (if available) from Sakornsakolpat et al., 2019 (GREx-genetically regulated expression, mQTL-methylation quantitative trait loci, Cod-coding association, Hi-C-chromatin interaction in human lung or IMR90 cell line, DHS-DNase hypersensitivity sites, GSet-genes identified by DEPICT, further details are available in the original publication); colocalization, probability shared causal variant between eQTL (GTEx) and COPD risk association (tissue: probability), only colocalization probability > 0.6 are listed. B) List of *Drosophila* genes and RNAi lines included in the screen. RNAi lines were ranked according to the natural logarithm of the ratio between the proportion of smurfs after candidate gene knockdown and luciferase RNAi control. Cutoff scale shown in Fig 1C was used to determine the effect of each RNAi. Based on this fine-grained ranking of individual RNAi lines, an overall rating was assigned to each gene and compared to human eqtl data (see also Fig 1A). Temperature column refers to the temperature the subsets of RNAi lines were screened at.

**Figure S1(Related to Figure 2).**
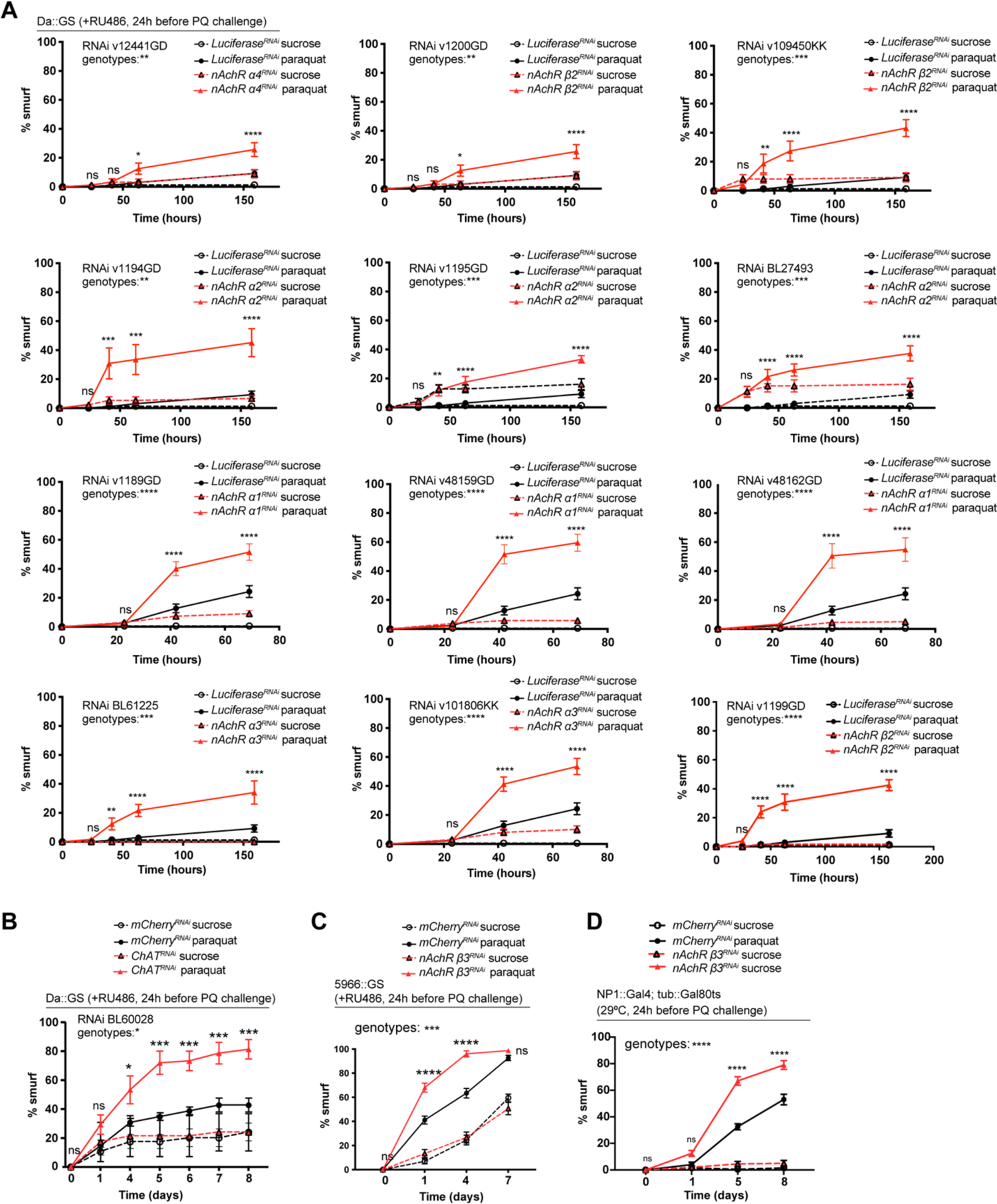
A) Barrier dysfunction assay after mCherry (control) or indicated nAchR subunit depletion for 24h with ubiquitous driver Da::GS. N=1. nAchR *α*4 (v12441GD): n=100 for Luciferase RNAi (control) on sucrose; n=125 animals for Luciferase RNAi on sucrose+paraquat; n=125 for nAchR *α*4 RNAi on sucrose; n=150 animals for nAchR *α*4 RNAi on sucrose+paraquat. nAchR *β*2(v1200GD): n=100 for Luciferase RNAi (control) on sucrose; n=125 animals for Luciferase RNAi on sucrose+paraquat; n=75 for nAchR *β*2 RNAi on sucrose; n=150 animals for nAchR *β*2 RNAi on sucrose+paraquat nAchR *β*2(v109450KK): n=100 for Luciferase RNAi (control) on sucrose; n=125 animals for Luciferase RNAi on sucrose+paraquat; n=125 for nAchR *β*2 RNAi on sucrose; n=150 animals for nAchR *β*2 RNAi on sucrose+paraquat nAchR *α*2 (v1194GD): n=100 for Luciferase RNAi (control) on sucrose; n=125 animals for Luciferase RNAi on sucrose+paraquat; n=125 for nAchR *α*2 RNAi on sucrose; n=150 animals for nAchR *α*2 RNAi on sucrose+paraquat. nAchR *α*2 (v1195GD): n=100 for Luciferase RNAi (control) on sucrose; n=125 animals for Luciferase RNAi on sucrose+paraquat; n=125 for nAchR *α*2 RNAi on sucrose; n=150 animals for nAchR *α*2 RNAi on sucrose+paraquat. nAchR *α*2 (BL27493): n=100 for Luciferase RNAi (control) on sucrose; n=125 animals for Luciferase RNAi on sucrose+paraquat; n=125 for nAchR *α*2 RNAi on sucrose; n=125 animals for nAchR *α*2 RNAi on sucrose+paraquat. nAchR *α*1 (v1189GD): n=175 for Luciferase RNAi (control) on sucrose; n=175 animals for Luciferase RNAi on sucrose+paraquat; n=175 for nAchR *α*1 RNAi on sucrose; n=175 animals for nAchR *α*1 RNAi on sucrose+paraquat. nAchR *α*1 (v48159GD): n=150 for Luciferase RNAi (control) on sucrose; n=150 animals for Luciferase RNAi on sucrose+paraquat; n=125 for nAchR *α*1 RNAi on sucrose; n=150 animals for nAchR *α*1 RNAi on sucrose+paraquat. nAchR *α*1 (v48162GD): n=1075 for Luciferase RNAi (control) on sucrose; n=175 animals for Luciferase RNAi on sucrose+paraquat; n=150 for nAchR *α*1 RNAi on sucrose; n=150 animals for nAchR *α*1 RNAi on sucrose+paraquat. nAchR *α*3 (BL61225): n=100 for Luciferase RNAi (control) on sucrose; n=125 animals for Luciferase RNAi on sucrose+paraquat; n=50 for nAchR *α*3 RNAi on sucrose; n=75 animals for nAchR *α*3 RNAi on sucrose+paraquat. nAchR *α*3 (v101806KK): n=175 for Luciferase RNAi (control) on sucrose; n=175 animals for Luciferase RNAi on sucrose+paraquat; n=175 for nAchR *α*3 RNAi on sucrose; n=175 animals for nAchR *α*3 RNAi on sucrose+paraquat. nAchR *β*2: n=100 for Luciferase RNAi (control) on sucrose; n=125 animals for Luciferase RNAi on sucrose+paraquat; n=75 for nAchR *β*2 RNAi on sucrose; n=100 animals for nAchR *β*2 RNAi on sucrose+paraquat. B) Barrier dysfunction assay after mCherry (control) or ChAT depletion for 24h with ubiquitous driver Da::GS. n=60 animals per genotype and condition; N=3. Two-way ANOVA followed by Šídák’s multiple comparisons test. C) Barrier dysfunction assay after mCherry (control) or nAchR *β*3 depletion for 24h with enterocyte-specific driver 5966::GS. nAchR *β*3: n=175 for mCherry RNAi (control) on sucrose; n=175 animals for mCherry RNAi on sucrose+paraquat; n=150 for nAchR *β*3 RNAi on sucrose; n=150 animals for nAchR *β*3 RNAi on sucrose+paraquat. N=3. Two-way ANOVA followed by Šídák’s multiple comparisons test. D) Barrier dysfunction assay after mCherry (control) or nAchR *β*3 depletion for 24h with enterocyte-specific driver NP1::Gal4, tub::Gal80^ts^ (NP1^ts^). n=200 animals per genotype and condition; N=3. Two-way ANOVA followed by Šídák’s multiple comparisons test. Data presented as mean ± SEM. ns, not significant, P > 0.05; *P ≤ 0.05; **P ≤ 0.01; ***P ≤ 0.001; ****P ≤ 0.0001. n: number of animals or midguts analyzed; N: number of independent experiments performed with similar results and a similar n

**Figure S2(Related to Figure 4).**
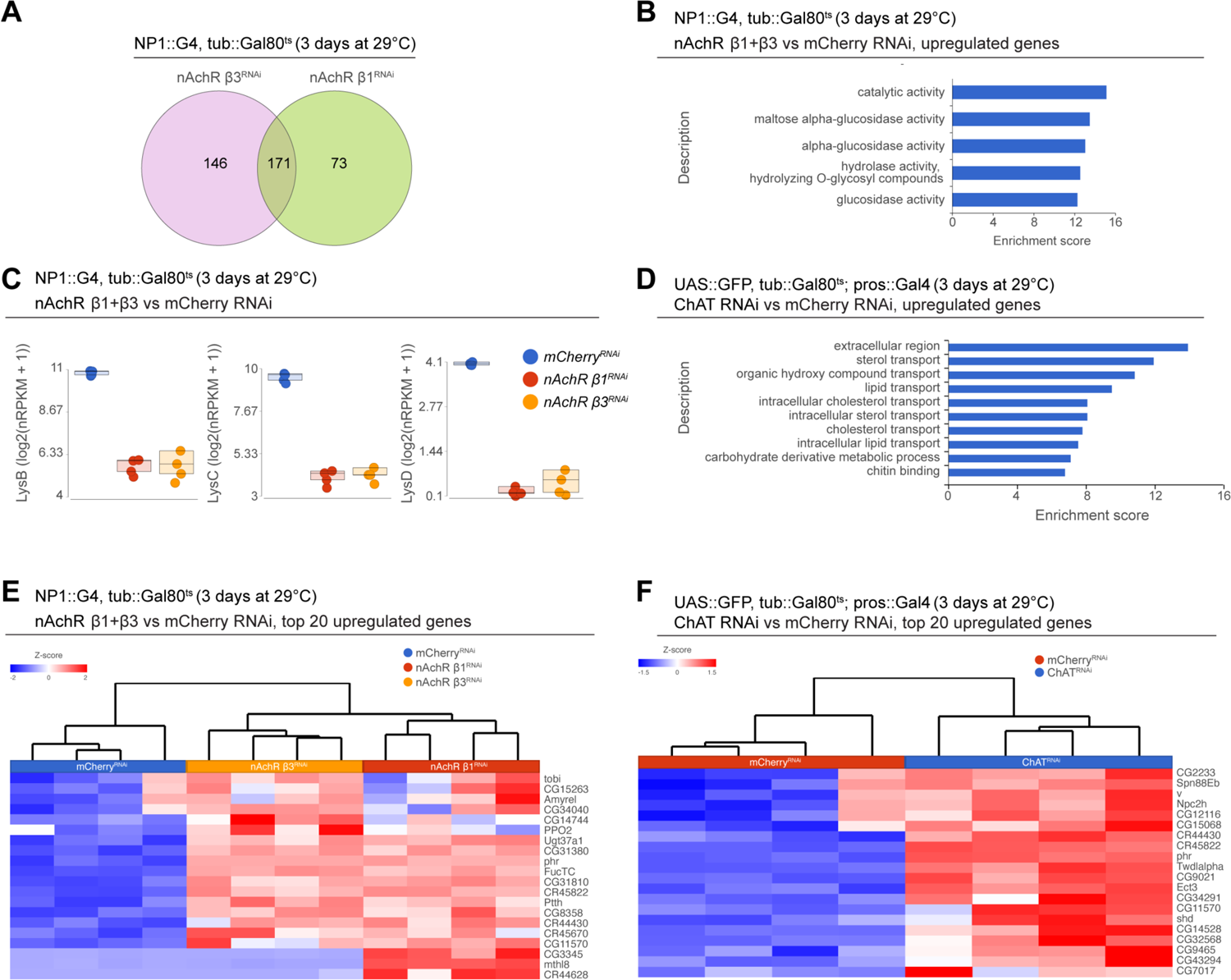
A) Overlap between differentially regulated genes after 3 days of nAchR *β*1 or *β*3 depletion in enterocytes with NP1^ts^. B) GO term enrichment of significantly upregulated genes after 3 days of nAchR *β*1 and *β*3 knockdown with NP1^ts^. C) Transcript levels of lysozyme family members in bulk RNAseq data set after 3 days of mCherry (control) or nAchR *β*1 and *β*3 subunit depletion in ECs. D) Gene set enrichment analysis of significantly upregulated genes after ChAT depletion in EEs. E) Heatmap of the top 20 upregulated differentially expressed genes after nAchR *β*1 and *β*3 knockdown in ECs for 3 days. (FDR<= 0.1; log2(fold change) > 2.5; 100% of samples have >= 1 reads)) F) Heatmap of top 20 upregulated differentially expressed genes after 3 days of ChAT depletion with RNAi in EEs under control of pros^ts^. (FDR<= 0.1; log2(fold change) > 1.97; 100% of samples have >= 1 reads)

**Figure S3(Related to Figure 5).**
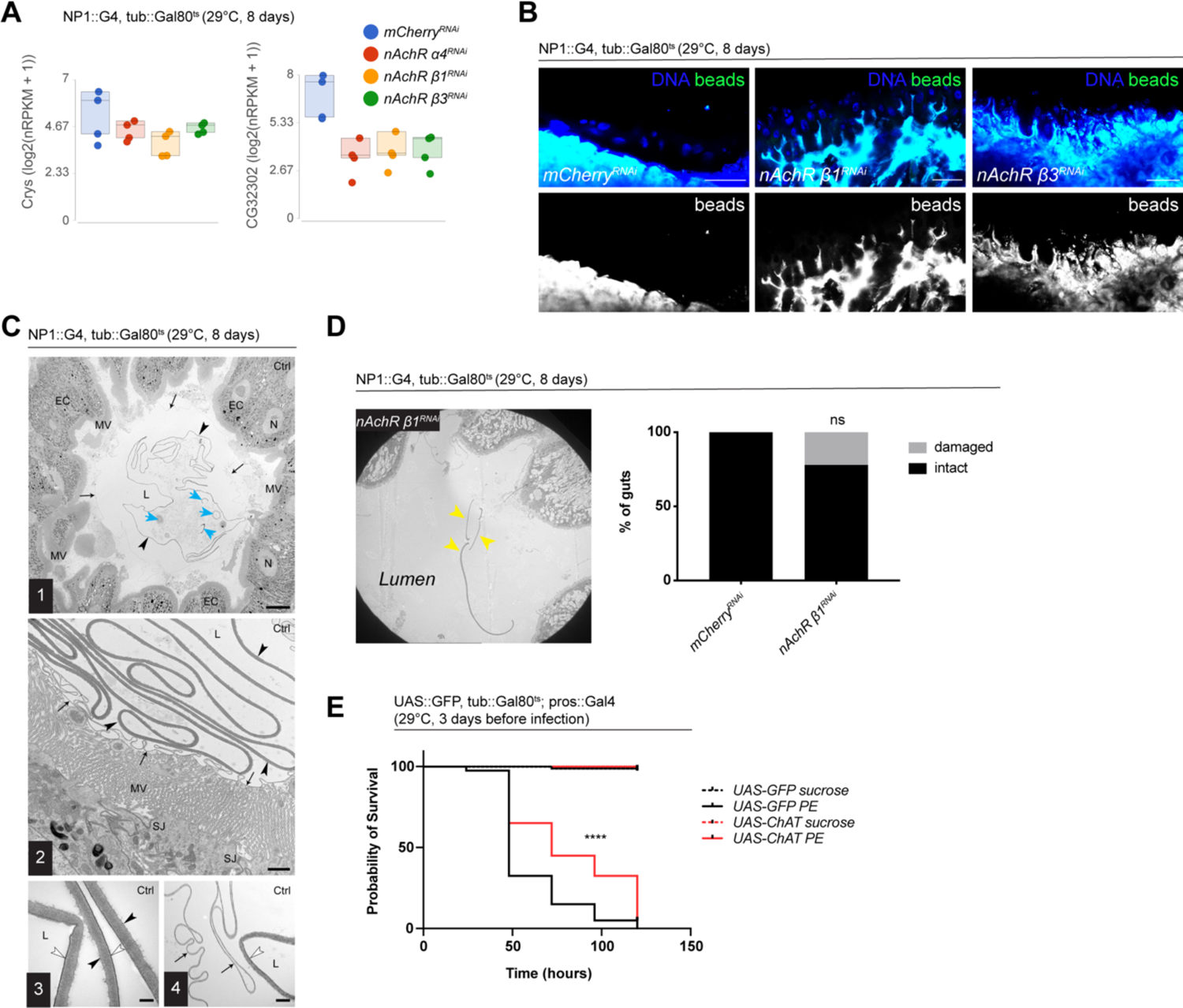
A) Transcript levels of two PM components in bulk RNAseq data set after 8 days of mCherry (control) or nAchR subunit depletion in ECs. B) Confocal immunofluorescence image of posterior midguts depleted for either mCherry (control), nAchR *β*1 or *β*3 for 8 days. Animals were fed green fluorescent beads to assess Peritrophic matrix (PM) integrity. DNA (blue) is labeled with Hoechst. n=10 guts. N=3. Scale bar 25μm. C) Overview of PM layers in posterior midgut (R4) of control animals. (1) The PM lies as an intact ring (black arrowheads) loosely in the gut lumen surrounded by an additional thin layer ring (small arrows). The PM encloses food remnants and short segments of material with a similar ultrastructure as the PM (blue arrows). (2) Detail of the PM layers: Thick layer (black arrowheads) and thin layer ring (small arrows) on top of microvilli of the enterocytes. Septate junctions (SJ) seal the intercellular spaces between the enterocytes at their apical edges. (3) Detail of the PM ultrastructure. The luminal surface (white arrowhead) is lined by an electron-dense layer of constant thickness. The abluminal surface is less electron-dense and slightly rough (black arrowheads). (4) Detail of the thin layer ring (black arrows) L, gut lumen. EC, enterocyte. MV, microvilli. N, nucleus. Scale bars: 10 µm (1), 1 µm (2), 200 nm (3), 500 nm (4). D) Example image of damaged thick PM layer, yellow arrowheads highlight PM fragments in the gut lumen. Quantification of thick PM layer integrity after 8 days of mCherry (control) or nAchR *β*1 knockdown in ECs. n=16; 18 midguts for mCherry or nAchR *β*1, respecitvely. N=1. Fisher’s exact test. E) Survival after 3 days of mCherry (ctrl) or ChAT overexpression in EEs followed by *Pseudomonas entomophila* infection. n=80 animals for UAS-GFP on sucrose or sucrose+PE; n=40 animals for UAS-ChAT on sucrose or sucrose+PE; N=3. Log Rank (Mantel-Cox) test. Data presented as mean ± SEM. ns, not significant, P > 0.05; *P ≤ 0.05; **P ≤ 0.01; ***P ≤ 0.001; ****P ≤ 0.0001. n: number of animals or midguts analyzed; N: number of independent experiments performed with similar results and a similar n.

**Figure S4(Related to Figure 6).**
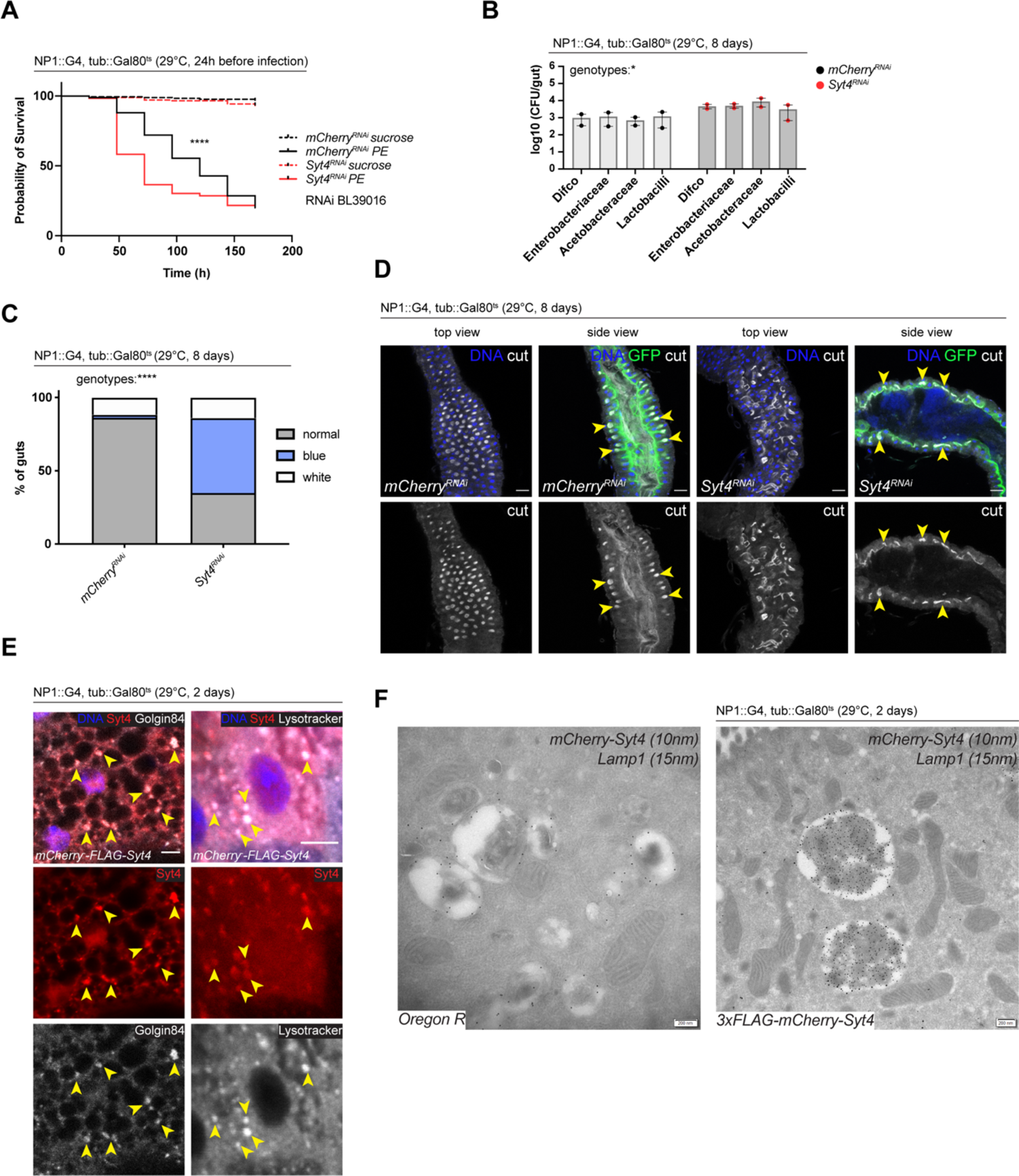
A) Survival after one day of mCherry (ctrl) or Syt4 depletion in ECs followed by *Pseudomonas entomophila* infection. n=175 animals per genotype and condition; N=3. Log Rank (Mantel-Cox) test. B) CFU of whole guts plated on selective growth media after 8 days of Syt4 depletion in ECs. 2 pooled independent experiments are shown. n=5 pooled animals per genotype and experiment. Two-way ANOVA. C) Analysis of gut compartmentalization visualized by feeding Bromphenol blue pH indicator (see Fig 5E) after 8 days of Syt4 knockdown in ECs. n=51 guts for mCherry (control), n=43 guts for Syt4 RNAi. 3 independent pooled experiments are shown. Chi square test. D) Confocal immunofluorescence images of posterior midguts depleted of either mCherry (control) or Syt4 in enterocytes for 8 days, stained with anti-cut antibody (white). Guts are expressing a GFP-brush border marker (green). DNA (blue) is labeled with Hoechst. Yellow arrowheads in side view panels highlight healthy, pocket-like (mCherry) and disrupted gastric units (Syt4-RNAi). n=10 guts. N=3. Scale bar 25μm. E) Confocal immunofluorescence image of posterior midguts overexpressing UAS-FLAG-mCherry-Syt4 (red) in enterocytes stained with anti-Golgin84 antibody or Lysotracker (white). DNA (blue) is labeled with Hoechst. Yellow arrowheads indicate overlap between Syt4-positive vesicles and Golgin84 or Lysotracker staining. n=10 guts. N=3. Scale bar 5μm. F) Immunogold electron microscopy image of Syt4 and Lamp1 co-staining in enterocytes of posterior midguts of either wildtype Oregon R flies or animals expressing UAS-FLAG-mCherry-Syt4 under control of NP1ts. Syt4 (detected with mCherry antibody 10nm gold particles) and Lamp1 (15nm gold particles) are colocalizing on multilamellar bodies in animals expressing the Syt4 construct. Oregon R samples are devoid of anti-mCherry antibody staining. n=5 guts. N=1. Scale bar 200nm. Data presented as mean ± SEM. ns, not significant, P > 0.05; *P ≤ 0.05; **P ≤ 0.01; ***P ≤ 0.001; ****P ≤ 0.0001. n: number of animals or midguts analyzed; N: number of independent experiments performed with similar results and a similar n.

